# An amygdalo-parabrachial pathway regulates pain perception and chronic pain

**DOI:** 10.1101/2020.01.10.902205

**Authors:** Charles Raver, Olivia Uddin, Yadong Ji, Ying Li, Nathan Cramer, Carleigh Jenne, Marisela Morales, Radi Masri, Asaf Keller

## Abstract

The parabrachial (PB) complex mediates both ascending nociceptive signaling and descending pain modulatory information in the affective/emotional pain pathway. We have recently reported that chronic pain is associated with amplified activity of PB neurons in a rat model of neuropathic pain. Here we demonstrate that similar activity amplification occurs in mice, and that this is related to suppressed inhibition to PB neurons from the central nucleus of the amygdala (CeA). Animals with pain after chronic constriction injury of the infraorbital nerve (CCI-Pain) displayed higher spontaneous and evoked activity in PB neurons, and a dramatic increase in after-discharges—responses that far outlast the stimulus—compared to controls. PB neurons in CCI-Pain animals showed a reduction in inhibitory, GABAergic inputs. We show that—in both rats and mice—PB contains few GABAergic neurons, and that most of its GABAergic inputs arise from CeA. These CeA GABA neurons express dynorphin, somatostatin and/or corticotropin releasing hormone. We find that the efficacy of this CeA-LPB pathway is suppressed in chronic pain. Further, optogenetically stimulating this pathway suppresses acute pain, and inhibiting it, in naïve animals, evokes pain behaviors. These findings demonstrate that the CeA-LPB pathway is critically involved in pain regulation, and in the pathogenesis of chronic pain.

**Significance Statement:** We describe a novel pathway, consisting of inhibition by dynorphin, somatostatin and corticotropin-releasing hormone expressing neurons in the central nucleus of the amygdala that project to the parabrachial nucleus (PB). We show that this pathway regulates the activity of pain-related neurons in PB, and that, in chronic pain, this inhibitory pathway is suppressed, and that this suppression is causally related to pain perception. We propose that this amygdalo-parabrachial pathway is a key regulator of both chronic and acute pain, and a novel target for pain relief.

## Introduction

Chronic pain is a public health concern that profoundly affects the quality of life of individuals worldwide (Dworkin et al., 2007; O’Connor and Dworkin, 2009; van Hecke et al., 2014). Chronic pain afflicts over 100 million Americans – more than heart disease, cancer, and diabetes combined. Pain costs the nation up to $650 billion a year in medical treatment and lost productivity (Institute of Medicine (US) Committee on Advancing Pain Research, 2011). Chronic pain is the most common complaint of patients in outpatient clinics (Upshur et al., 2006). Because the underlying pathophysiology of chronic pain is largely unknown, current pharmaceutical or surgical therapies provide minimal relief to chronic pain patients (Meyer-Rosberg et al., 2001; Dworkin et al., 2013). The ultimate goal of this research program is to rectify this deficiency. Improved understanding of the neurophysiological, cellular, and molecular mechanisms that contribute to neuropathic pain is critically needed to develop more effective therapeutic strategies.

Most previous attempts to pinpoint the pathophysiology of chronic pain have focused on the lateral, sensory-discriminative system, including the somatosensory thalamus and cortex. This is despite the inability of this approach to spawn more effective therapies. There is growing evidence that therapies targeting the motivational-cognitive dimensions of pain may be more promising (Auvray et al., 2010). Indeed, there is increasing evidence that the negative affective, cognitive, and psychosocial state of chronic pain is universal in different chronic pain states (Gustin et al., 2011). Therefore, understanding the role of the affective-motivational pathways in chronic pain may lead to innovative therapies to treat these widespread conditions.

A key structure for encoding the affective component of pain is the parabrachial complex (PB). PB comprises the Kölliker-Fuse nucleus and the lateral and medial parabrachial nuclei. It is located at the midbrain-pons junction and derives its name from its proximity to the superior cerebellar peduncle. PB has extensive, often reciprocal connections with brainstem, medulla and forebrain structures, and it plays key roles in functions such as satiety and appetite, sleep and arousal, cardiovascular function, and fluid homeostasis (Hajnal et al., 2009; Martelli et al., 2013; Davern, 2014). Importantly, PB plays a particularly prominent role in pain processing (Gauriau and Bernard, 2002; Roeder et al., 2016, 2016; Uddin et al., 2018; Chiang et al., 2019).

PB receives dense inputs from lamina I nociceptive spinal neurons, a pain-related projection – far denser than the spinothalamic pathway (Spike et al., 2003; Polgár et al., 2010). As a result, PB neurons can respond robustly—and preferentially—to noxious stimuli (Gauriau and Bernard, 2002; Uddin et al., 2018). PB projects, often reciprocally, to several regions linked to pain and affect, including the periaqueductal gray, rostroventral medulla, thalamus, amygdala and zona incerta (Bianchi et al., 1998; Roeder et al., 2016). Thus, PB appears to serve as a key nexus for pain and its affective perception.

PB is involved also in chronic pain. Matsumoto et al. (1996) reported that PB neuronal activity is increased in a rat model of arthritic pain. Expression of immediate early genes in PB increases after chronic constriction injury (CCI) of the sciatic nerve in rats (Jergova et al., 2008). We demonstrated that chronic pain after CCI of the infraorbital nerve (CCI-ION) is associated with amplified activity of PB neurons (Uddin et al., 2018). This amplified activity was expressed as a dramatic increase in after-discharges, the excessively prolonged neuronal responses that far outlast a sensory stimulus. Because these after-discharges are likely causally related to chronic pain (Laird and Bennett, 1993; Asada et al., 1996; Okubo et al., 2013), it is important to understand the mechanisms leading to their generation, and the circuitry involved in their pathophysiology.

Here, we describe a novel pathway—the CeA-LPB pathway—that potently inhibits PB neurons. This inhibition regulates PB responses to nociceptive inputs, and dysregulation of this inhibition leads to amplified activity of PB neurons, and to chronic pain.

## Materials and Methods

We adhered to accepted standards for rigorous study design and reporting to maximize the reproducibility and translational potential of our findings as described by Landis et al. (2012) and in ARRIVE (Animal Research: Reporting In Vivo Experiments). In line with NIH recommendations for scientific rigor, we performed an a priori power analysis to estimate required sample sizes (Landis et al., 2012).

### Subjects

All procedures adhered to Animal Welfare Act regulations, Public Health Service guidelines, and approved by the University of Maryland School of Medicine Animal Care and Use Committee. We studied male and female adult rats and mice, unless otherwise noted. We studied 14 Sprague-Dawley rats (acquired from Envigo Indianapolis, IN), 11 corticotropin releasing hormone-Cre (CRH-Cre) rats (generously provided by R. Messing, University of Texas, Austin) bred on a Wistar background, 43 wild-type Wistar rats (from our breeding colony, founding breeders acquired from Envigo), 3 GAD2-GFP mice (generously provided by A. Puche, University of Maryland, Baltimore) bred on a B6CBAF1/J (Jackson Laboratories, Bar Harbor, ME) background, 21 GAD2-IRES-Cre mice (Jackson Laboratories), and 15 C57Bl6/J background mice (from our breeding colony, founding breeders from Jackson Laboratories).

### Recovery surgical procedures

#### Induction of chronic orofacial pain

We used a rodent model of neuropathic pain, evoked by chronic constriction of the infraorbital nerve (CCI-ION) (Vos et al., 1994; Benoist et al., 1999; Okubo et al., 2013; Akintola et al., 2017; Castro et al., 2017a). We first anesthetized animals with isoflurane, followed by ketamine/xylazine (i.p.). We made intraoral incisions along the roof of the mouth next to left cheek, beginning distal to the first molar. We freed the infraorbital nerve from surrounding connective tissue before loosely tying it with silk thread (4–0), 1–2 mm from its emergence from the infraorbital foramen. After the surgery, we monitored the animals daily as they recovered for 5–7 days in their home cage.

#### Viral construct and anatomical tracer injections, iontophoresis, and cannula implants

We placed animals, deeply anesthetized with isoflurane, in a stereotaxic frame and created a small craniotomy over the region of interest, targeting those regions under stereotaxic guidance. For delivery of viral constructs to the CeA, we injected 0.8-1 µL of the viral construct at a rate of 50 nL/min, using glass pipettes (40-60 µm tip diameter), coupled to a Hamilton^®^ syringe controlled by a motorized pump. The pipette was left in place for 10 minutes before being slowly retracted over 5-10 mins. We targeted the CeA using the following coordinates: in rats (AP −2.2 mm and ML 4.3 mm, relative to Bregma, and DV −6.7 mm relative to dural surface) and in mice (AP −1.1 mm and ML 2.6 mm, relative to Bregma, and DV −4.0 mm relative to dural surface).

For delivery of retrograde anatomical tracers to the PB of rats and mice, we used either cholera toxin subunit B (CTB, List Biological Labs) or Fluoro-Gold™ (FG; Fluorochrome, LLC, Denver), as noted in Results. We injected 1 µL of 0.5% CTB (List Biological Labs, Campbell, CA) in saline through glass pipettes (tip diameter 40 µm) at a rate of 50 nL/min. For the delivery of FG, we iontophoretically injected 3% tracer dissolved in 0.1M cacodylate buffer using glass pipettes (10-15 µm tips). We delivered pulses of 5 µA (positive polarity, 5-7 sec duty cycle, 30 min). We targeted PB using the transverse sinus and the following coordinates: in rats (AP −8.6 to −9.0 mm and ML 2.0 mm, relative to Bregma, and DV −5.3 mm relative to dural surface) and in mice (AP −4.6 mm and ML 1.2 mm, relative to Bregma, and DV −3.2 mm relative to dural surface).

To pharmacologically or optogenetically manipulate PB, we implanted either stainless steel guide cannulas (26G, Plastics One, Roanoke, VA), or fiber optic cannulas (200 µm core, 0.37 NA, RWD Life Science, San Diego) using coordinates listed above for rats. Throughout the post-implantation recovery period, we handled animals regularly to acclimate them to removal of the cannula’s protective cap and connection to the internal cannula or fiber optic patch cable.

### Behavioral assessment of pain and hyperalgesia

#### Mechanical sensitivity

To assess tactile sensitivity, we held animals loosely without restraint on the experimenter’s arm (rats) or hand (mice), while von Frey filaments (North Coast Medical, Gilroy, CA) of varying forces were applied to the buccal region. We tested each animal bilaterally, and a response was defined as an active withdrawal of the head from the probing filament. We used the up-down method to determine withdrawal thresholds, as described previously (Dixon, 1965; Chaplan et al., 1994; Akintola et al., 2017). We used the same approach to assess tactile responses on the plantar surface of the hindpaws, in animals standing on a metal grid. We compared grouped data with Mann-Whitney U ranked-sum tests or Friedman’s non-parametric repeated measures tests as noted (see Results).

#### Dynamic allodynia

To assess dynamic mechanical sensitivity, we performed a modified brush test (adapted from Cheng et al., 2017). Briefly, we placed animals in a clear Plexiglass chamber on a raised wire platform to provide unrestricted access to the hindpaw. One round consisted of three brush stimuli applied to the lateral plantar surface of the hindpaw at intervals of 10s. We performed three rounds, alternating left and right hindpaws with 3 mins between rounds. The stimuli consisted of a light stroke from proximal to distal end of the plantar surface, using a clean #4 paintbrush trimmed flat. We counted the number of responses out of 9 total stimulus applications for each hindpaw. A response consisted of flicking, licking, or a complete lifting of the hindpaw from the floor, whereas we did not count only partial movement of the paw or limb that did not result in the paw lifting from the platform.

#### Grimace scale

To assess ongoing pain, we analyzed facial grimace behavior (Langford et al., 2010; Sotocinal et al., 2011; Akintola et al., 2017, 2019). We placed animals in a square Plexiglas™ chamber (6×8 inches for rats, 3×5 inches for mice) with two opaque sides and two transparent sides on a raised wire platform. For all experiments, except optogenetic stimulation, we recorded each animal for 30 mins, with cameras facing each of the transparent sides of the chamber. For the optogenetics experiments, we placed animals in a modified Plexiglass™ chamber with three transparent sides, and recorded throughout the mechanical sensitivity tests.

We scored facial expressions via a semi-automated procedure using the “Face Finder” application (Sotocinal et al., 2011) generously gifted by J. Mogil. We screened, labelled, scrambled and scored face images with the experimenter blinded to the treatment group and identity of each image using a custom Matlab (MathWorks, Naticks, MA) script. The grimace scale quantifies changes in four action units: orbital tightening, nose-cheek bulge, whisker tightening and ear position. We selected ten screenshots for each animal or each time point (during within-subject and repeated measures tests), and on each image, each action unit was given a score of 0, 1, or 2, as previously described (Langford et al., 2010; Sotocinal et al., 2011; Akintola et al., 2017, 2019). We calculated mean grimace scale scores as the average score across all the action units.

### *In vivo* electrophysiology

#### Surgical preparation

We lightly anesthetized (Level III-2, as defined by Friedberg et al., 1999) C57Bl6/J mice with 20% urethane. We placed mice in a stereotaxic frame, with body heat maintenance, and made a small craniotomy over the recording site to target PB (AP −4.0 to −4.6 mm and ML 1.2 to 1.5 mm, relative to Bregma, and DV −3.0 to −3.5 mm, relative to dural surface). Post-hoc histological analysis and small electrolytic lesions confirmed all recording sites; cells falling outside of PB were excluded. To identify recording sites, electrolytic lesions were made at the end of a recording session. We sectioned the fixed brain tissue into 80 μm-thick coronal sections that were stained with cresyl violet.

#### Electrophysiological recording

Using platinum-iridium recording electrodes (2–4 MΩ) produced in our laboratory, we recorded from PB ipsilateral to CCI-ION. We isolated units responsive to noxious cutaneous stimuli, to dermatomes in both the head and body, and digitized the waveforms using a Plexon system (Plexon Inc., Dallas, TX). Upon encountering a neuron responsive to noxious cutaneous stimulation, we allowed the neuron to resume baseline firing rate before recording spontaneous activity for three minutes, after which we recorded neuronal responses to noxious stimuli. We applied mechanical stimuli within the V2 dermatome, or to the plantar surface of the hind-paw. Mechanical stimulation was produced with calibrated electronic forceps. We applied five repetitions each of mechanical stimulus, alternating between ipsilateral and contralateral receptive fields. When the neuron recorded resumed firing at its baseline rate, we applied stimuli with at least 8 s between each application. If neurons exhibited after-discharges, we extended the inter-stimulus interval to capture the entire after-discharge duration.

#### Electrophysiology data analysis

We sorted neurons using Offline Sorter (Plexon Inc.) using dual thresholds and principal component analysis. We subsequently generated autocorrelograms in NeuroExplorer (Plexon Inc.) to confirm that each recording was of a single unit. We analyzed responses to tactile stimuli using custom Matlab routines, written to calculate the integral of force applied by the forceps, the firing rate during stimulus application, and the spontaneous firing rate. Evoked responses were computed and expressed as evoked firing rate normalized to spontaneous firing rate, divided by the stimulus force integral (Castro et al., 2017b).

We defined after-discharges—periods of sustained activity that outlast a stimulus presentation (Okubo et al., 2013; Uddin et al., 2018) as PSTH bins in which activity exceeded the 99% confidence interval for a period lasting at least 500 ms after stimulus offset.

### *In vitro* electrophysiology

#### Surgical preparation of animals

At least three weeks prior to *in vitro* electrophysiological recordings, we injected CeA of GAD-Cre mice (21-30 days old) with channelrhodopsin viral constructs, as described above, AAV-EF1a-DIO-hChR2(H134R)-EYFP (UNC Vector Core, Chapel Hill, NC). Animals recovered for 5-7 days from injection of the viral construct before being assessed for baseline mechanical sensitivity. Animals underwent CCI-ION surgery (as described above) or sham surgery, and were allowed to recover for an additional two weeks. Just before electrophysiological recordings, we measured mechanical sensitivity thresholds to confirm the presence or absence of CCI-ION pain.

#### *In vitro* recordings

We anesthetized animals with ketamine/xylazine, removed their brains, and prepared horizontal slices (300-µm-thick) containing PB, following the method described by Ting et al. (2014). For recordings, we placed slices in a submersion chamber and continually perfused (2 ml/min) with ACSF containing (in mM): 119 NaCl, 2.5 KCl, 1.2 NaH_2_PO_4_, 2.4 NaHCO_3_, 12.5 glucose, 2 MgSO_4_·7H_2_O, and 2 CaCl_2_·2H_2_O.

We obtained whole-cell patch-clamp recordings, in voltage-clamp mode, through pipettes containing (in mM): 130 cesium methanesulfonate, 10 HEPES, 1 magnesium chloride, 2.5 ATP-Mg, 0.5 EGTA, and 0.2 GTP-Tris. For recordings in bridge mode, we replaced cesium methanesulfonate with potassium gluconate (120 mM) and potassium chloride (10 mM). Impedance of patch electrodes was 4 – 6 MΩ. Series resistance (<40 MΩ was monitored throughout the recording, and recordings were discarded if series resistance changed by >20%. All recordings were obtained at room temperature.

To optically activate ChR2, we collimated blue light through a water-immersion 40X microscope objective to achieve whole-field illumination. Light source was a single wavelength (470 nm) LED system (CoolLED pE-100, Scientifica, Clarksburg, NJ), controlled through a TTL signal.

### Pharmacological and optogenetic manipulation of PB

#### Inhibition of PB

To assess whether inhibiting PB can alleviate CCI-ION pain, we first implanted wild-type Wistar rats with a single guide cannula above the right PB, as described above. Two weeks later, we recorded baseline facial and hindpaw mechanical withdrawal thresholds and RGS scores. Animals then underwent CCI-ION surgery, and, 2 months later, we obtained mechanical and spontaneous ongoing pain scores. We randomly assigned animals to a saline or muscimol group, and infused 50 µL of 50 µM muscimol, or saline, at 2.5 µL/min, into PB via internal cannula temporarily inserted into the chronic guide cannula. We immediately tested mechanical withdrawal thresholds and RGS scores following drug infusion. After 1 week to allow for drug washout, we switched the treatment groups (drug/saline) and animals were retested. We confirmed cannula implantation locations by *post hoc* histological analysis.

#### Disinhibition of the PB

To assess whether disinhibiting PB can evoke behavioral metrics of pain, we implanted wild-type Wistar rats with a single cannula in the right PB, similar to the muscimol experiments above. Animals recovered for two weeks after implantation before testing baseline hindpaw mechanical withdrawal thresholds. We randomly assigned animals to saline or gabazine groups., and infused 50 µL of 200 nM gabazine, or saline, at 2.5 µL/min into PB. We immediately tested hindpaw mechanical withdrawal thresholds. We normalized each animal’s post-drug behavioral score to its baseline score.

#### Optogenetic activation of the CeA-PB terminals

To determine if selectively activating the CeA-PB pathway can alter pain sensation, we injected wild-type Wistar or CRH-Cre rats with channelrhodopsin viral constructs, and implanted fiber optic cannulas, as described above. We injected AAV-hSyn-hChR2(H134R)-mCherry (or AAV-hSyn-hChR2(H134R)-EYFP) and AAV-EF1a-DIO-hChR2(H134R)-EYFP into the right CeA of wild-type Wistar and CRH-Cre rats, respectively. In all animals, we implanted a single fiber optic cannula over the right PB. Animals recovered for 9 weeks before behavioral testing. Although fluorescent reporter protein expression can be observed in PB starting around 6 weeks post-injection, expression appears to peak and stabilize around 8-9 weeks.

We tested all fiber optic cannulas before implantation and calibrated the laser light source at the beginning of each testing session to deliver 4 mW of 470 nm light at the fiber optic terminus. We stimulated CeA-PB terminals with short trains of 5 stimuli (4 ms duration/stimulus, 10 Hz) from a 470 nm laser controlled by TTL pulse from Master-8 pulse generator (A.M.P.I., Jerusalem). The experimenter triggered each train of light manually, with a 5 s delay, to ensure application of mechanical and dynamic hindpaw stimuli could be time locked with light delivery. During all testing sessions, we connected animals to the fiber optic patch cable and held constant other behavioral cues with the light stimulation portion of the testing session.

After the 9-week recovery, we collected baseline mechanical sensitivity, dynamic mechanical sensitivity, and RGS scores. In naïve animals, we performed dynamic mechanical test with animals connected to the fiber optic patch cable without light delivery, waited for a 5 min recovery period, and repeated the test with delivery of light time locked to each hindpaw stimulus. We then returned animals to their home cage and allowed 90-120 min rest period. We then performed the mechanical withdrawal test in the same manner – hindpaw stimuli applied but no light delivery, 5 min rest period, hindpaw stimuli applied with concurrent light delivery. For the acute formalin pain test, we repeated this same testing paradigm after injecting 50 µL of 5% formalin (Sigma-Aldrich), injected s.q. into the dorsal surface of the left hindpaw (contralateral to injection and cannula implantation). We waited approximately 90 minutes to allow the initial pain response—lifting, licking, and guarding of the hindpaw—to subside before starting the mechanical sensitivity tests. We videotaped every session to gather RGS scores with and without optogenetic activation of the CeA-PB. We collected RGS scores in this manner to avoid prolonged activation of this pathway in the absence of a pain stimulus, as well as the confounding effects of prolonged blue light exposure (Tyssowski and Gray, 2019).

### Anatomical, immunohistochemical and neurochemical techniques

#### Tissue preparation

After a post-injection recovery period, specific to the tracer and species (see below), we deeply anesthetized animals with ketamine/xylazine, or urethane, and perfused them transcardially with phosphate buffered saline (PBS), followed by 4% paraformaldehyde (Sigma-Aldrich). We removed brains and immersed them in fixative overnight at 4°C. Using a vibrating microtome (50 µm sections) or a cryostat (14-16 µm sections), we collected coronal sections through CeA and PB.

#### Anterograde viral tracing

We injected CeA of GAD-Cre mice and CRH-Cre rats with AAV-EF1a-DIO-hChR2-EYFP viral constructs (see above). After at least 3 weeks for mice and 9 weeks for rats, we perfused, collected, and sectioned tissue. We rinsed free-floating sections (50 µm) of the injection site (CeA) and projection site (PB) several times in PBS, and counterstained with DAPI (100 ng/mL for 5 min). We then mounted and cover-slipped tissue with a non-fluorescing hydrophilic mounting media.

#### Identification of inhibitory neurons in PB

We collected brains from formalin-perfused Wistar rats, sectioned (14 µm thick) and mounted frozen sections directly onto Superfrost Plus slides (Fisher Scientific, Hampton, NH). We processed sections for mRNA transcripts encoding the vesicular GABA transporter (VGAT) using RNAscope*^®^* (Advanced Cell Diagnostics, Newark, CA). According to the manufacturer’s directions, we treated sections with heat, protease digestion, and hybridization of the target probes with Opal 520 fluorophore (PerkinElmer, Waltham, MA). Following hybridization of the fluorophore via RNAscope, we treated all sections for NeuN immunoreactivity using rabbit anti-NeuN antibody (abcam, Cambridge, MA; ab128886; 1:500). We blocked sections for 30 mins at room temperature in 2% normal donkey serum with 0.2% Triton X-100, followed by incubation in rabbit anti-NeuN for 24 hours at 4°C. We washed sections twice in PBS, followed by incubation with Cy3 conjugated donkey anti-rabbit (JacksonImmuno; #711-165-152; 1:300) for 1 hr at room temperature. We washed sections twice in PBS, counterstained with DAPI, and cover-slipped with ProLong™ Gold mounting medium (Invitrogen, Carlsbad, CA). Each batch included sections processed with negative and positive control probes to ensure specificity of the target probes.

Following RNAscope and NeuN immuno-labelling, we quantified VGAT expressing LPB neurons from images photographed with a Leica Microsystems (Wetzlar, Germany) TCS SP8 confocal microscope. We held excitation and detection parameters constant across all sections. We imaged full sections using a 5X objective, and LPB and surrounding structures using 40X oil immersion objective. We reconstructed sections using Leica LAS X Navigator tiling software. Finally, we analyzed sections using StereoInvestigator (MBF Bioscience, Williston, VT) after drawing boundaries for LPB (Paxinos and Watson, 2013). In each section, we identified and quantified VGAT and NeuN-positive cells by dividing LPB in to 100 µm x 100 µm grids and sampling 25% of each grid. We analyzed the left and right LPB of four sections spaced 280 µm apart from each animal.

#### Retrograde immunohistochemical identification of CeA-PB neurons

We injected PB of GAD-GFP mice and Sprague-Dawley rats with CTB (see above). Between 7 to 14 days post injection, we perfused, collected, and sectioned tissue. We processed free-floating sections (50 µm) for double-label immunohistochemistry with antibodies against CTB (goat anti-CTB; List Biological Labs, Inc.; product #703; 1:20,000) and against calbindin (rabbit anti-calbindin D-28k; Swant, Switzerland product #CB38; 1:10,000). We incubated sections for 48-72 hours at 4°C with anti-CTB and anti-calbindin antibodies in 4% normal donkey serum with 0.1% Triton X-100, before washing several times in PBS at room temperature. We then incubated sections for 1 hr at room temperature with Cy3 conjugated donkey anti-rabbit (JacksonImmuno, West Grove, PA; product #711-165-152; 1:1000) and Alexa488 conjugated donkey anti-goat (JacksonImmuno; product #705-545-147; 1:1000). After several washes in PBS, we counterstained with DAPI, mounted and cover-slipped tissue slides.

#### Retrograde identification of CeA-PB neurons using in situ hybridization (ISH)

We injected PB of Sprague-Dawley and Wistar rats with FG (see above). After 14-21 days post injection, we perfused, collected, and sectioned tissue. We used a combination of in situ hybridization and immunolabeling. We incubated free-floating CeA coronal sections (16 µm) for 2 h at 30 °C with rabbit anti-FG antibody (Millipore; AB153I; 1:200) in DEPC-treated phosphate buffer (PB) with 0.5% Triton X-100 supplemented with RNasin (40Ul/µl stock, 5 ml/ml of buffer, Promega). We rinsed sections, (3 × 10 min) in DEPC-treated PB and incubated in biotinylated donkey anti-rabbit antibody (Vector Laboratories; 1:200) for 1 h at 30 °C. We rinsed sections with DEPC-treated PB and then transferred to 4% PFA. We then rinsed sections with DEPC-treated PB, incubated for 10 min in PB containing 0.5% Triton X-100, rinsed with PB, treated with 0.2N HCl for 10 min, rinsed with PB, and then acetylated in 0.25% acetic anhydride in 0.1M triethanolamine, pH 8.0, for 10 min. We rinsed sections with PB, and post-fixed with 4% PFA for 10 min. Before hybridization and after a final rinse with PB, we incubated the sections in hybridization buffer for 2 h at 55 °C (50% formamide, 10% dextran sulfate, 5 × Denhardt’s solution, 0.62M NaCl, 50mM DTT, 10mM EDTA, 20mM PIPES, pH 6.8, 0.2% SDS, 250 mg/ml salmon sperm DNA, 250 g/ml tRNA). Sections were hybridized for 16 h at 55 °C in hybridization buffer containing [35S]- and [33P]- labelled single-stranded antisense CRF (nucleotides 1-1093; GenBank accession number: X03036) probes at 10^7^ c.p.m./ml. We treated sections with 4 mg/ml RNase A at 37 °C for 1 h and washed with 1 × saline-sodium citrate, 50% formamide at 55 °C for 1 h and with 0.1 × saline-sodium citrate at 68 °C for 1 h. We then rinsed sections with PB and incubated for 1 h at room temperature in avidin-biotinylated horseradish peroxidase (Vector Laboratories; ABC kit; 1:100). Sections were rinsed, and the peroxidase reaction was developed with 0.05% 3,30-diaminobenzidine (DAB) and 0.03% H2O2. We mounted sections on coated slides and dipped slides in Ilford K.5 nuclear tract emulsion (Polysciences, 1:1 dilution in double-distilled water) and exposed in the dark at 4 °C for 4 weeks before development.

#### Retrograde identification and quantification of CeA-PB neurons using RNAscope^®^

We injected PB of Wistar rats with FG (see above). We perfused, collected, and sectioned tissue (14 µm) and mounted frozen sections directly onto Superfrost Plus slides (Fisher Scientific). We processed sections for mRNA transcripts encoding somatostatin, prodynorphin, and corticotropin releasing hormone (CRH) using RNAscope*^®^*, as described above. We labelled all sections for somatostatin, and alternated labeling adjacent sections for either dynorphin or CRH. According to the manufacturer’s directions, we treated sections with heat, protease digestion, and hybridization of the target probes with fluorophores, Opal 520 and Opal 650 (PerkinElmer, Waltham, MA). Following hybridization of fluorophores via RNAscope, we treated all sections for FG immunoreactivity using rabbit anti-FG antibodies (MilliporeSigma, Burlington, MA; AB153-I; 1:500). We blocked sections for 30 mins at room temperature in 2% normal donkey serum with 0.2% Triton X-100, followed by incubation in rabbit anti-FG for 24 hours at 4 °C. We washed sections twice in PBS, followed by incubation with Cy3 conjugated donkey anti-rabbit (JacksonImmuno; #711-165-152; 1:300) for 1 hr at room temperature. We washed sections twice in PBS, counterstained with DAPI, and cover-slipped with ProLong™ Gold mounting medium (Invitrogen, Carlsbad, CA). Each batch included sections processed with negative and positive control probes to ensure specificity of the target probes.

Following RNAscope and FG immuno-labelling, we quantified the neurochemical phenotype of CeA-PB neurons by observing and photographing sections with a Leica Microsystems (Wetzlar, Germany) TCS SP8 confocal microscope. We held excitation and detection parameters constant across all sections. We imaged full sections using a 5X objective, and CeA and surrounding structures using 20X oil immersion objective.

We reconstructed sections using Leica LAS X Navigator tiling software. Finally, we analyzed sections using Neurolucida (MBF Bioscience, Williston, VT), thresholding the FG signal to reduce non-specific immunolabelling, and manually drawing boundaries for CeA (Paxinos and Watson, 2013). We manually identified and quantified FG-positive neurons, followed by identifying those neurons that co-labelled for somatostatin, dynorphin, and CRH. We collected every fifth section through CeA and counted every neuron within CeA. We included only sections with at least 20 FG-positive neurons to control for variations in sectioning angle, edge effects, and deposition or uptake of FG at the injection site. From each of the 5 animals analyzed, we included approximately 20 sections (see Results) for this quantification. We compared distributions of somatostatin labelled FG-positive neurons between the two assays (*i.e.* RNAscope for somatostatin and dynorphin versus somatostatin and CRH) and found no differences.

### Statistical analysis

We analyzed group data using GraphPad Prism version 8 for Mac (GraphPad Software, La Jolla CA). Data are presented, unless otherwise noted, as median values ± 95% confidence intervals (95% CI). We used non-parametric Mann-Whitney U tests, Wilcoxon matched-pairs signed rank tests, and Friedman’s repeated measures tests, as noted below (see Results).

## Results

### CCI-ION results in signs of pain

As we and others described previously (Bennett and Xie, 1988; Akintola et al., 2017). CCI-ION resulted in pain-like behaviors in mice, assessed 3 weeks after surgery. Thresholds for withdrawal from mechanical stimuli in CCI-ION mice were more than twofold lower (median of 1.2 g; 0.3 to 2.2 g, 95% CI) compared to shams (2.8 g; 1.8 to 4.2 g; p=0.009, Mann Whitney *U*= 2; Fig. 1A). This indicates that CCI-ION resulted in persistent hyperalgesia.

**Figure 1.**
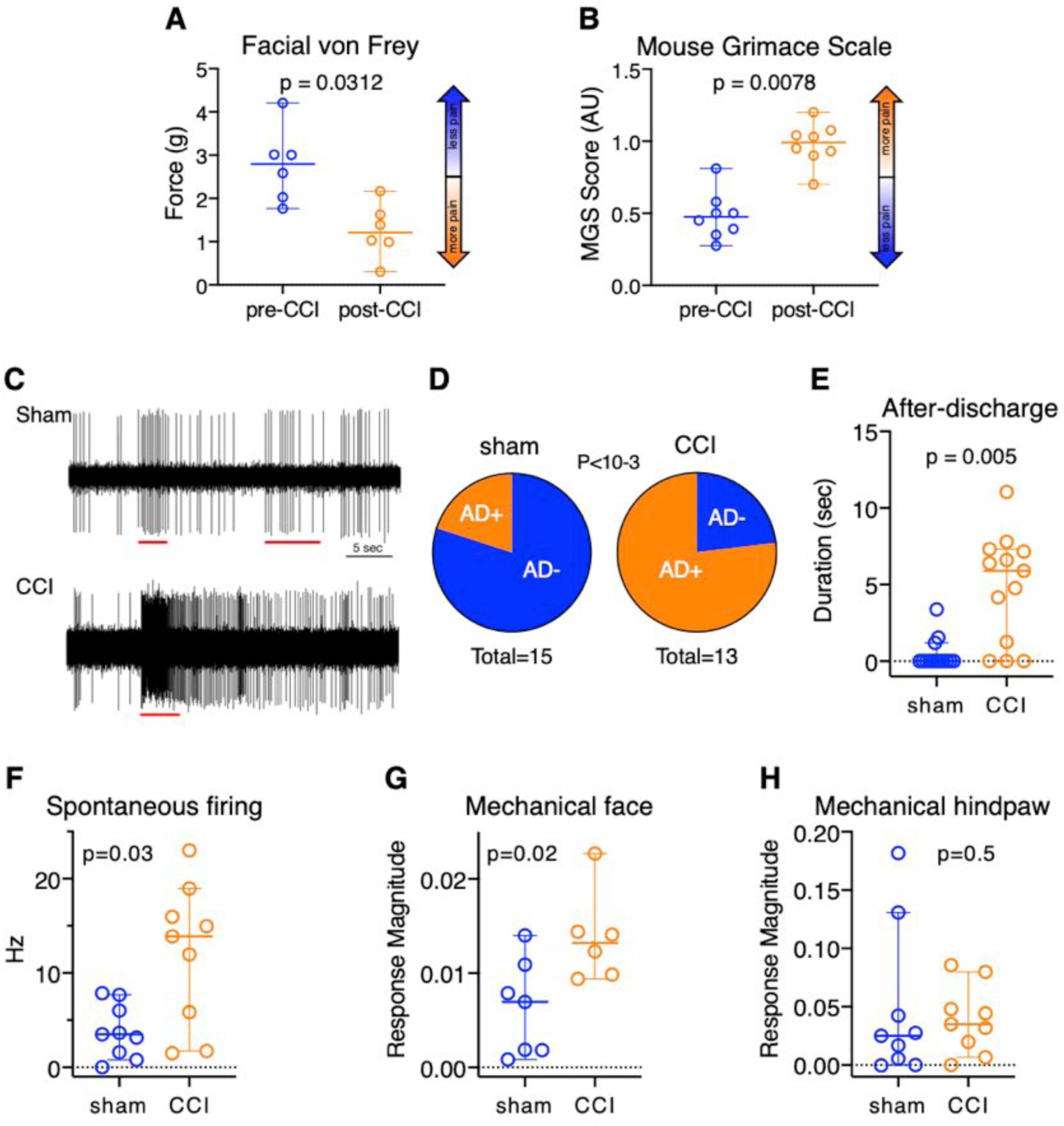
CCI-ION causes behavioral signs of pain and amplification of neuronal activity in the parabrachial nucleus. **A-H**, Data are presented as medians and 95% confidence intervals. In CCI-ION mice, mechanical withdrawal thresholds are lower (**A**), and mouse grimace scale scores are higher (**B**). **C**, Sample traces of extracellular recordings from a sham and a CCI-ION animal, with red bars depicting timing of the application of mechanical stimuli. In PB neurons recorded from CCI-ION mice, the duration of after-discharges is longer (**E**), and the proportion of tactile responsive PB neurons displaying after-discharges is higher after CCI-ION (**D**). In CCI-ION mice, PB neuron spontaneous firing rates are higher (**F**), and magnitudes of responses to mechanical stimuli applied to the face (**G**) and hindpaw (**H**) are higher.

We have previously reported that the mouse grimace score is a reliable and sensitive metric for the assessment of ongoing pain in mice with CCI-ION (Akintola et al., 2017). Consistent with this, CCI-ION mice had grimace scores that were more than twofold-higher than those in sham mice (median = 0.99, 0.70 to 1.20 95%CI *vs* 0.475, 0.275 to 0.810; p=0.0003, Mann Whitney *U* =1; Fig. 1B).

These findings indicate that mice with CCI-ION exhibited signs of both ongoing (“spontaneous”) pain, as well as mechanical hypersensitivity.

### PB neuron activity is amplified after CCI-ION

We have previously demonstrated that parabrachial activity is amplified in rats with chronic pain, induced by CCI-ION (Uddin et al., 2018). Here we demonstrate that this pathological phenomenon occurs also in mice. We also show that—like in the rat— CCI-ION is associated with amplified responses to stimuli applied both to the face and to the hindlimb.

Fig. 1C depicts representative extracellular spikes recorded from isolated PB neurons from an anesthetized, sham-operated animal (1C, top) and from a CCI-ION mouse (1C, bottom). Activity evoked in response to face pinch, applied through calibrated forceps, was 8-fold higher in the CCI-ION neuron, compared to the sham. Significantly, the PB neuron from the CCI-ION mouse showed pronounced after-discharges (ADs)— responses that far outlasted the stimuli—that were of longer duration and frequency, compared to those in the control neuron. In this neuron, ADs lasted more than 40 sec.

PB neurons recorded from CCI-ION mice were four times as likely to exhibit after-discharges, with 77% of tactile responsive neurons from CCI animals and 20% of neurons from sham animals displaying ADs (p <10^-3^, binomial test, Fig. 1D). AD duration was longer in CCI-ION mice (n = 13, median = 5.9 sec, 95% CI = 0 to 7.3 sec) than in sham-operated mice (n = 14, median = 0 secs, 95% CI = 0 to 1.2; Mann-Whitney U = 27.50, p = 0.005; Cohen’s *d* = 1.7 (“large” effect size); Fig. 1E).

We quantified responses of PB neurons *during* noxious application of calibrated forceps to the face or hindpaws. Neuronal response magnitudes are expressed as arbitrary units, computed as frequency (in Hz) divided by stimulus magnitude (integral of force applied). The magnitude of response to stimulation of the face, but not the hindpaw, was higher in CCI-ION animals, compared to controls (Fig. 1G-H). Neurons recorded from mice with CCI responded to facial stimuli with a median magnitude of 0.013 units (95% CI = 0.009 to 0.023, n = 6), whereas neurons recorded from sham injured animals responded with a median magnitude of 0.007 units (95% CI = 0.001 to 0.014, n = 7; Mann-Whitney *U* = 5; p = 0.02; Fig. 1G; Cohen’s *d* = 1.5 (“large” effect size)). In contrast, neurons from both conditions responded similarly to hindpaw stimuli; sham animals (n = 9) had a median response magnitude of 0.025 units (95% CI = 0.000 to 0.131) and CCI-ION cells had a median response of 0.035 units (95% CI = 0.007 to 0.080; Mann-Whitney *U* = 32.5, p = 0.5; Fig. 1H).

Spontaneous firing rates of PB neurons in CCI-ION mice were higher than those in control animals. Neurons from CCI-ION mice (n = 9) had a median firing rate of 13.9 Hz (95% CI = 1.7 to 19.0 Hz), whereas neurons from sham animals (n = 9) fired at median firing rate of 3.5 Hz (95% CI = 0.8-7.7 Hz; Mann-Whitney *U* = 16, p = 0.03; Fig. 1F; Cohen’s *d* = 1.4 (“large” effect size)).

These data indicate that—as we previously reported in rats (Uddin et al., 2018)—CCI-ION results in amplification of PB neuronal activity. In mice, CCI-ION manifested as an increase in spontaneous firing rates and in larger responses to mechanical stimuli. Most prominent was the increase in the magnitude of ADs in animals with pain after CCI.

### Inhibitory inputs to PB are reduced after CCI-ION

We and others have previously shown that amplified neuronal activity in chronic pain can result from maladaptive disinhibition of CNS structures (reviewed in Masri and Keller, 2012; Prescott, 2015). To test if disinhibition is associated with the amplified activity of PB neurons after CCI-ION, we recorded from PB neurons in acute slices. Figure 2A depicts traces of whole-cell recordings from PB neurons from CCI-ION and control mice. Miniature inhibitory postsynaptic currents (mIPSCs) appear as inward currents because of the high [Cl^2-^] in the patch pipette (see Methods). The frequency of these events is approximately 50% lower in the example from the CCI-ION animal, compared to the control. For group comparisons, we first averaged data from all neurons recorded from a particular mouse, and used these averages as our individual samples (that is, sample size=number of mice). This analysis revealed that the frequency of mIPSCs in CCI-ION mice was approximately half that in controls (Cohen’s *d* = 1.4). Median mIPSC frequency in PB cells from CCI-ION mice (n = 12 mice) was 1.6 Hz (95% CI: 0.4-2.0 Hz), compared with 2.8 Hz (95% CI = 1.4-4.1; Mann-Whitney *U* = 11, p = 0.02; Fig. 2B) in sham animals (n = 6). In contrast, the amplitudes of mIPSCs in CCI-ION mice was indistinguishable from that in controls. Neurons from CCI-ION mice (n = 12 mice) had a median mIPSC amplitude of 23.5 pA (95% CI = 17.3-40.0 pA), and neurons from sham animals (n = 6) had at median amplitude of 29.8 pA (95% CI = 10.1-39.1 pA; Mann-Whitney *U* = 35, p = 0.9; Fig. 2C).

**Figure 2.**
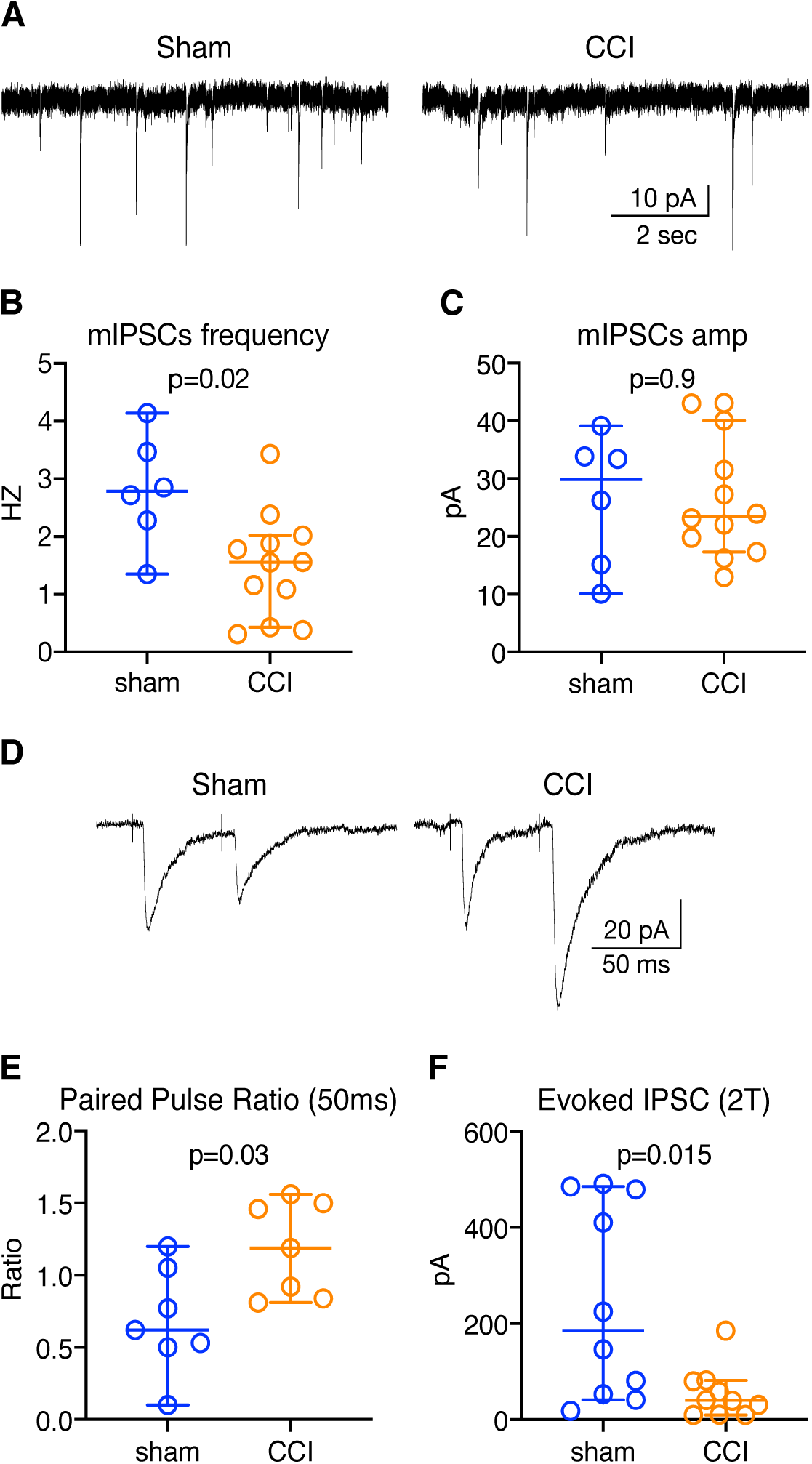
CCI-ION attenuates inhibitory inputs to the lateral parabrachial (LPB) from the central nucleus of the amygdala (CeA). Data are presented as medians and 95% confidence intervals. ***A***, Example recordings of miniature inhibitory postsynaptic currents (mIPSCs) from PB neurons from sham and CCI mice. ***B***, mIPSCs frequency is lower in CCI-ION mice, whereas their amplitudes remain indistinguishable from controls (C). ***D***, Example traces of IPSCs evoked in LPB neurons by optogenetically activating inhibitory CeA afferents with pairs of light stimuli. ***E***, Paired pulse ratios (50 msec) are higher, consistent with a reduction in synaptic efficacy. ***F***, Amplitude of IPSC evoked by stimulating CeA afferents at twice stimulus threshold are lower in CCI-ION mice.

Changes in the amplitudes of synaptic potentials might be masked by parallel changes in input resistance of the recorded neurons. However, there was no difference (p = 0.9; Mann-Whitney U = 225) in resistance of neurons from CCI-ION mice (median = 771 MΩ; 95%CI = 631 to 983 MΩ), compared to controls (median = 691 MΩ; 95%CI = 617 to 1056 MΩ). In current clamp recordings, there was no difference (p = 0.19; Mann-Whitney U=176) in resting membrane potential of neurons from CCI-ION mice (median = −68 mV; 95%CI = −69 to −67), compared to controls (median = −69 mV; 95%CI = −70 to −67 mV).

### Source of inhibition

PB is reported to contain only a small number of inhibitory neurons (Guthmann et al., 1998; Yokota et al., 2007). Consistent with these findings, examination of sections through PB of GAD2-GFP mouse revealed that PB, and the lateral parabrachial nucleus in particular, contain only a small number of GAD-GFP neurons (Fig. 3A). In contrast, other regions, such as the cerebellum, contain a high density of these neurons.

**Figure 3.**
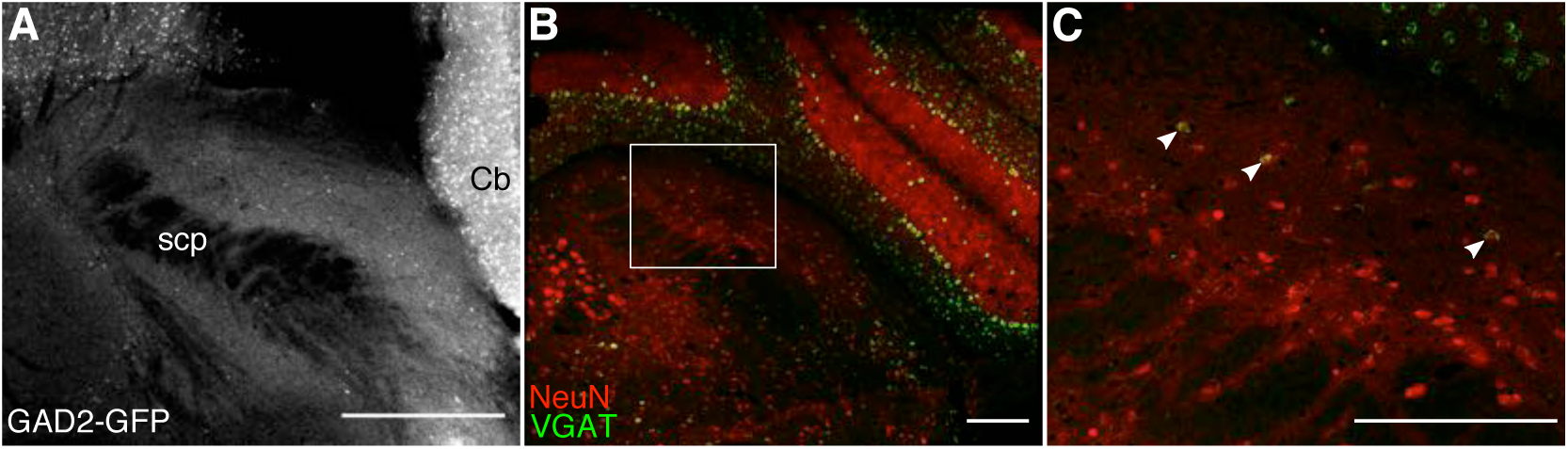
Parabrachial nuclei contain few inhibitory, GABAergic interneurons. ***A***, A representative image of coronal sections from GAD2-GFP mice depicts few GABAergic neurons (bright white puncta) in PB. ***B-C***, Few LPB neurons express VGAT. Representative images of coronal sections through the PB of Wistar rats showing immunoreactivity for NeuN (red) and expression of VGAT mRNA (green). Square inset shows a portion of LPB enlarged in panel ***C*** (white arrows denote example VGAT neurons). Scale bars equal 250 µm.

Similar to mice, sections through the PB of Wistar rats reveal a relatively a small proportion of PB neurons express VGAT (Fig. 3B-C). The lateral and ventral portions of the PB contain the densest collection of these GABAergic neurons, and a small number of isolated VGAT neurons appear throughout the PB. We estimated the proportion of GABAergic LPB neurons by quantifying the number of NeuN immuno-positive cells that express VGAT mRNA in non-adjacent coronal sections (see Methods). We calculated the proportion for each individual section, sampled at 280 µm-intervals from throughout the rostro-caudal extent of LPB. The median percentage of VGAT neurons was 12% (95% CI = 9 to 15%; n = 8 sections, 1 male and 1 female rat). The relative paucity of GABAergic neurons in LPB suggests that major inhibitory inputs to PB are extrinsic to this nuclear complex.

### Mouse anatomy

#### Retrograde tracing

To identify the inhibitory inputs that might be involved in disinhibition of PB following CCI-ION, we used retrograde and anterograde anatomical tracing, in both the rat and mouse. Injection of a retrograde neuronal tracer, cholera toxin subunit B (CTB), into the right PB of GAD2-GFP mice (n = 3), revealed a high density of retrogradely-labeled neurons in the ipsilateral central nucleus of the amygdala (CeA). We found lower density of retrograde labeling in the zona incerta and subthalamic nucleus, in the paraventricular and lateral hypothalamus, and in the insular cortex, all ipsilateral to the injection site. Figures 4A-C show coronal sections through the CeA, demonstrating that retrogradely-labeled neurons occupy much of the central nucleus.

**Figure 4.**
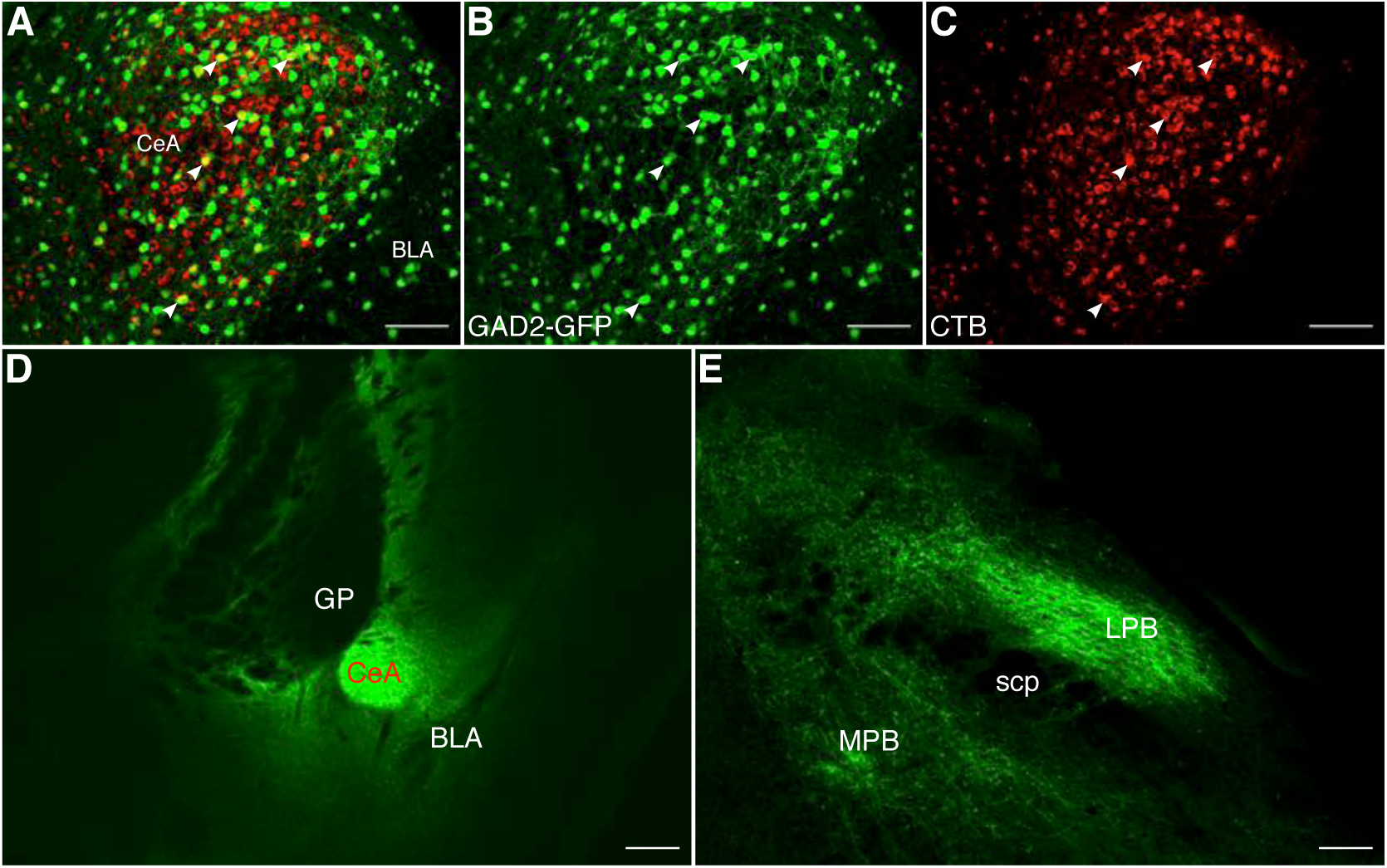
Retrograde and anterograde tracing in mice reveal that CeA provides dense inhibitory input to PB. ***A-C***, Representative images of coronal sections through the CeA of GAD2-GFP mice that received injections of CTB in PB. Merged signals (***A)*** from GAD2-GFP (***B***) and CTB (***C***) reveal CeA-PB projection neurons, identified with white arrows. ***D-E***, Representative images of coronal sections through CeA and LPB, respectively, from GAD2-Cre mice that received injection of a Cre-dependent viral construct (AAV-DIO-hChR2-eYFP) into CeA. Abundant axons expressing eYFP can be seen in LPB (***E***). Scale bars equal 100 µm (***A-C***) and 250 µm (***D-E***).

Because CeA is thought to be composed almost exclusively of GABAergic neurons (Duvarci and Pare, 2014), these findings indicate that inhibitory neurons in the mouse CeA provide dense inhibitory inputs to PB. Consistent with this conclusion, many neurons in CeA that were retrogradely labeled following CTB injection in PB were GAD2-positive (Fig. 4A-C arrows). We further characterize the chemical identity of these CeA-LPB neurons below.

#### Anterograde tracing

We confirmed the existence of an CeA-LPB pathway using an anterograde tracing approach. We injected into the right CeA of GAD-Cre mice (n = 3) the viral construct, AAV-EF1a-DIO-hChR2-EYFP (this construct was used for optogenetic manipulation studies described below). Images of sections through the injection site demonstrate that EYFP expression is restricted primarily to CeA (Fig. 4D). Sections through PB reveal a high density of labelled axons in PB (Fig. 4E). These projections from the CeA were notably dense in the lateral regions of PB, ipsilateral to the injection site. These findings confirm the existence of a dense CeA-LPB inhibitory pathway in the mouse.

### Rat Anatomy

#### Retrograde tracing

To determine if a similar CeA-LPB pathway exists in the rat we first used the retrograde neuronal tracer, cholera toxin subunit B (CTB). Figure 5A depicts a coronal section through PB of an adult rat injected with CTB, revealing an injection site covering much of the PB region. We made similar injections in the right PB of 6 adult male Sprague-Dawley rats. These injections resulted in dense, retrograde labeling of somata in the ipsilateral CeA (Fig. 5B-C). Similar to the mouse, we found a lower density of retrogradely labeled neurons in the zona incerta, the paraventricular and lateral hypothalamus, and the insular cortex, ipsilateral to the injection.

**Figure 5.**
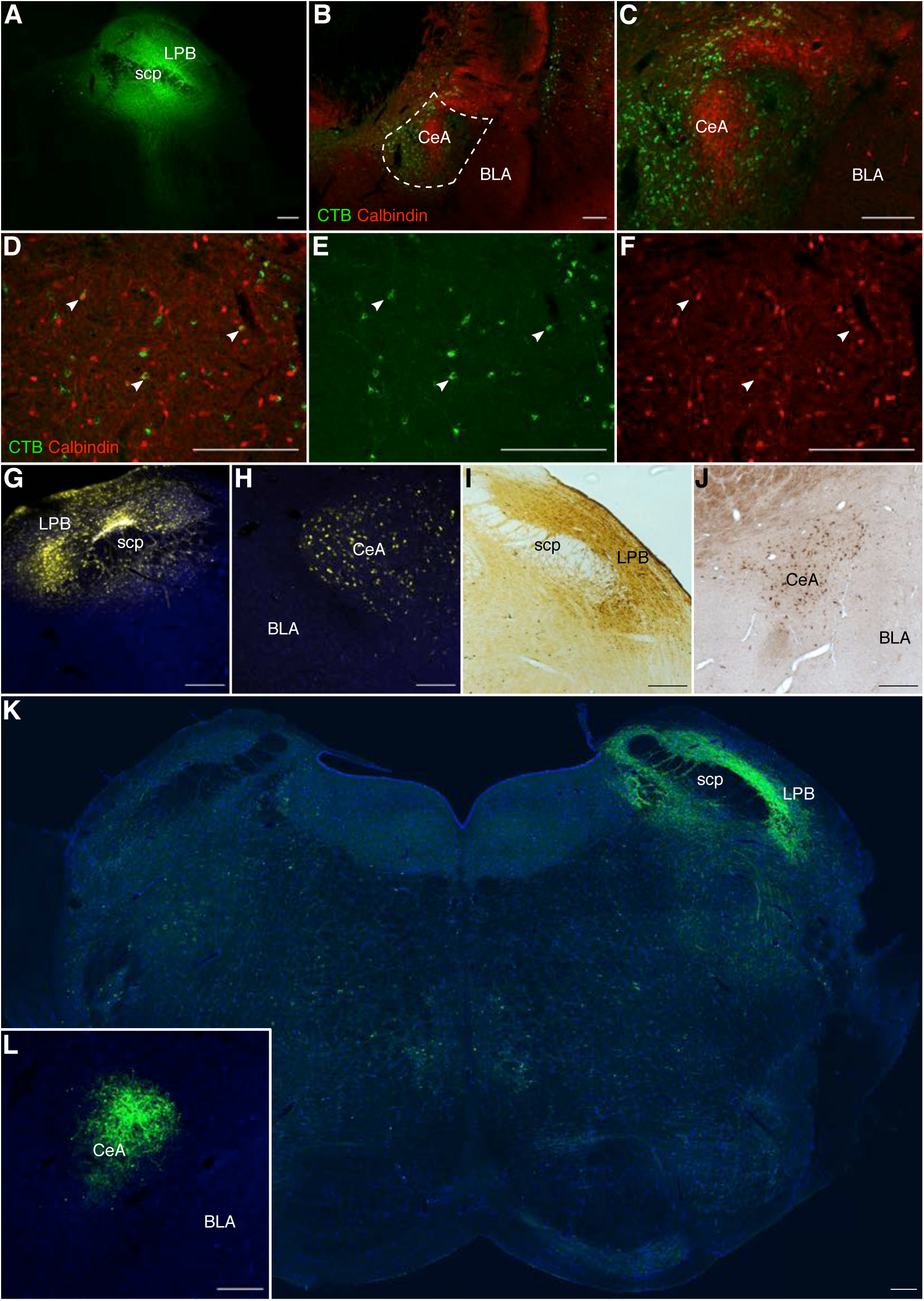
Retrograde and anterograde tracing in rats reveals that CeA provides robust inhibitory input to PB. ***A-F***, Representative images of coronal sections through PB and CeA of rats that received injections of CTB. ***A***, Dense deposition of CTB (immunoreactivity in green) at the injection site in PB. ***B-C***, Numerous retrogradely labelled neurons are seen in CeA in low and high magnification images of merged immunoreactivity signals from CTB (green) and calbindin (red). High magnification image of merged (***D***), CTB (***E***) and calbindin (***F***) signals reveal sparse co-labelling of individual neurons (white arrows). ***G-J***, Representative images of coronal sections through PB and CeA of rats that received discrete ionotophoretic injections of Fluoro-Gold™ (FG) into PB. Restricting FG depositions primarily to the LPB (***G*** and ***I***) revealed that the densest input to LPB arises from CeA (***H*** and ***J***). FG depositions and labelled neurons were identified by their fluorescent signal (yellow in ***G-H***) and immunoreactivity (brown reaction product ***I-J***). ***K-L***, Representative images of coronal sections through LPB and CeA, respectively, of CRH-Cre rats that received injection of a Cre-dependent viral construct (AAV-DIO-hChR2-eYFP) into CeA. Scale bars equal 250 µm.

To determine the phenotype of these retrogradely labeled neurons, we double-labeled the sections with antibodies for markers of GABAergic neurons (see Methods). Figure 5B-F shows examples of these retrogradely labeled neurons immunoreactive for the calcium binding protein, calbindin. Note that a large proportion of CTB-positive CeA neurons did not label for calbindin (Fig. 5 C-D), suggesting that they represent other classes of GABA neurons, a possibility we explore below. These data suggest that, like in the mouse, the CeA-LPB pathway in the rat consists of a dense projection of a heterogenous population of GABAergic CeA neurons.

We confirmed these findings using a different retrograde tracer, FluoroGold™ (FG). By making discrete iontophoretic injections of FG, we were better able to restrict our injections to those portions of the PB that have been previously implicated in processing and relaying nociceptive information (Cechetto et al., 1985; Roeder et al., 2016; Chiang et al., 2019). Further, while the CTB experiments in the rat were performed in male Sprague-Dawley rats, FG tracing was done in Sprague-Dawley and Wistar rats of both sexes. We observed no difference in the pattern of CeA-LPB projection between male and female animals, nor between the two strains of rat.

Figure 5 shows coronal sections through PB, demonstrating injection sites revealed with reflected (Fig. 5G) and transmitted light (Fig. 5I) microscopy. In contrast to our CTB injections above, injections of FG were restricted primarily to the lateral parabrachial (LPB). We observed strong deposition of FG throughout the LPB, with little dissipation into the medial parabrachial or other nearby nuclei, such as the Köllicker-Fuse or the locus coeruleus. We made similar injections into the right LPB of Sprague-Dawley rats (n = 3 of each sex) and bilaterally into the LPB of Wistar rats (n = 3 of each sex).

Sections through the amygdala reveal a pattern of labelled somata similar to that observed after CTB injections (Fig. 5 H&J). Neurons retrogradely labelled with FG were distributed throughout the ipsilateral CeA, but did not extend into the basolateral or medial nuclei of the amygdala. Below we describe our quantification of these projections, and identification of their chemical phenotypes. Consistent with our injections of CTB, we observed a small number of FG+ cells in the paraventricular and lateral hypothalamus, in the zona incerta, and insular cortex, all ipsilateral to the injection site. We observed no differences in either the pattern of distribution or laterality of these retrogradely identified cells between sex or animal strain.

#### Anterograde tracing

We confirmed the retrograde data in the rat with the use of anterograde viral tracing. We expanded the study by using a transgenic rat that expresses Cre exclusively in corticotropin releasing hormone (CRH) neurons (Pomrenze et al., 2015). This allowed us to restrict the expression of the viral constructs to CeA, as the expression of CRH in the amygdala is limited to this structure (Pomrenze et al., 2015 and Figs. 5&6).

**Figure 6.**
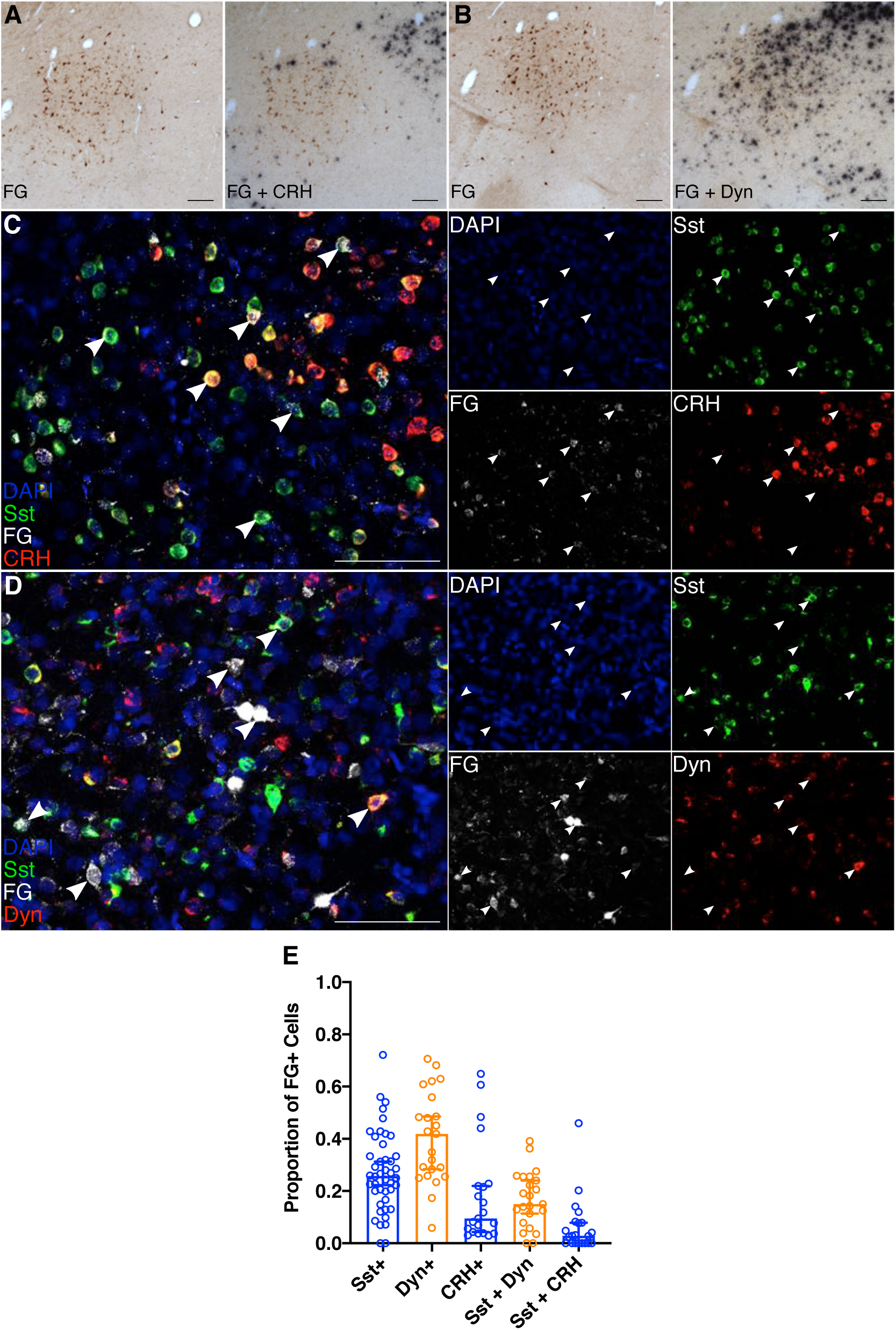
Neurochemical identification of CeA-LPB neurons reveals a heterogenous population expressing dynorphin, somatostatin and CRH mRNAs. ***A-B***, Representative images of coronal sections through CeA reveal limited expression of CRH mRNA (***A***) but robust expression of dynorphin mRNA (***B***) in CeA-LPB neurons. Left panels depict FG immunoreactive cells (brown reaction product) prior to deposition of silver grains via ***in situ*** hybridization (ISH). Right panels depict the same section after labelling for CRH or dynorphin mRNA (black reaction product), in ***A*** and ***B***, respectively. ***C-D***, Representative images of coronal sections through CeA demonstrate that CeA-LPB neurons can express somatostatin, CRH, or dynorphin mRNA. Large panels depict representative high-magnification images of merged signal from DAPI (pan-cellular stain; blue), somatostatin (green), FG (white), and CRH (red) in ***C*** or dynorphin (red) in ***D***. White arrows identify neurons that are FG+, and that co-label with one or both of the assayed mRNAs. Small panels depict the individual signals seen in the respective large panel, with white arrows retained to confirm neurochemical phenotype. ***E***, Median and 95% CI for the proportion of FG+ neurons that co-labeled for one or both assayed mRNAs. Scale bars equal 250 µm.

Injection of the viral construct AAV-DIO-hChR2-EYFP into the CeA of CRH-Cre rats (n=2 of each sex) resulted in highly localized expression within the CeA (Fig. 5L). These injections resulted in dense labeling of axons in the PB, primarily in ipsilateral LPB, but also included MPB (Fig. 5K). We observed no difference in the laterality or the distribution of projections between male and female rats.

These data suggest that the CeA-LPB pathway in the rat includes a dense projection from inhibitory neurons in the CeA to the LPB. Given the evidence from retrograde and anterograde anatomical approaches in both rats and mice, CeA is well positioned to provide a robust source of inhibitory control over nociceptive information processing in the lateral parabrachial.

### CCI-Pain results in suppressed CeA-PB inputs

The dense GABAergic, inhibitory projection from CeA to PB suggests that this pathway is involved in the reduced inhibition of PB neurons after CCI-ION. We tested this hypothesis by comparing CeA-LPB inputs in control and CCI-ION mice. In live slices through PB, we recorded inhibitory postsynaptic currents (IPSCs) from LPB neurons, evoked by optogenetic stimulation of CeA afferents to PB (Fig. 2D-F). We accomplished this by expressing ChR2 selectively in GABAergic neurons, by injecting Cre-dependent viral constructs in CeA of GAD2-CRE mice (see Methods, and “Source of Inhibition”).

Brief (0.3 msec) pulses of light evoked robust IPSCs in LPB neurons (Fig. 2D). To compare the efficacy of these evoked IPSCs in different neurons, we evoked pairs of stimuli, separated by 50 msec, and computed the paired pulse ratio (PPR), where a higher ratio is associated with “weaker” synapses, and a lower ratio with “stronger” synapses (Zucker, 1989; Kim and Alger, 2001; Sanabria et al., 2004). Because PPR is sensitive to stimulus intensity, we calibrated the optical stimulus for each neuron—by varying laser intensity—to obtain a response that was at twice the threshold stimulus intensity. For each neuron, we calculated PPR as the mean of the second response divided by the mean of the first (Kim and Alger, 2001), and then averaged the PPRs computed for all neurons recorded from the same animal. These averages were then used for statistical comparisons.

PPR recorded from neurons in CCI mice was nearly twice as large compared to that in control animals. Neurons from CCI-ION mice (n = 7 mice) had a median PPR of 1.19 (95% CI: 0.81-1.56), whereas neurons from sham animals (n = 7) had a median PPR of 0.62 (95% CI = 0.10-1.20; Mann-Whitney *U* = 7, p = 0.03; Fig. 3E; Cohen’s *d* =1.4).

The amplitudes of the first IPSCs, evoked at twice stimulus threshold, was nearly 5-fold smaller in CCI-ION mice, compared to controls (Cohen’s *d* =1.3). Neurons from CCI-ION mice (n = 10 mice) had a median evoked IPSC amplitude of 40.1 pA (95% CI: 9.9-81.8 pA), whereas neurons from sham animals (n = 10) had at median amplitude of 185.6 pA (95% CI = 41.2-482.2 pA; Mann-Whitney *U* = 18, p = 0.015; Fig. 2F).

These data suggest that inhibitory inputs from CeA to LPB are reduced in animals with chronic pain, compared to sham-operated controls. The reductions in mIPSC frequency and evoked IPSC amplitude, the increase in PPR, with no change in mIPSC amplitude, all point to pre-synaptic changes in this CeA-LPB circuitry. These electrophysiological data, coupled with our anatomical evidence, suggest that the amygdalo-parabrachial pathway may play a causal role in the development of neuropathic pain after CCI.

### Neurochemical identity of CeA-PB neurons

To further characterize the neurochemical phenotype of CeA-PB neurons we used a combination of *in situ* hybridization (ISH) and RNAscope. These revealed that CeA-LPB neurons are a heterogenous population, including neurons expressing corticotropin releasing hormone (CRH), somatostatin, and dynorphin; A subset of these CeA-LPB neurons expressed both somatostatin and dynorphin mRNA, or somatostatin and CRH mRNA.

Figure 5 (G&I) shows coronal sections through the PB of FG-injected rats. Only animals in which the injection site was restricted to LPB were used for further processing via ISH (n = 2 Sprague-Dawley rats and 3 Wistar rats) or RNAscope (n = 2 female, 3 male Wistar rats).

Figure 6 (A-B) depicts coronal sections through CeA of FG injected rats that were processed for ISH, and visualized with a combination of bright-field and epifluorescence microscopy. FG-positive neurons occupy much of the CeA. FG-positive neurons expressing mRNA for either CRH or dynorphin are depicted in Figures 6A and 6B, respectively. A small proportion of CeA-LPB neurons expressed CRH, whereas a substantial proportion of the FG labelled-positive population of CeA neurons was positive for dynorphin mRNA expression.

Figure 6 (C-D) shows coronal sections through the CeA of FG injected rats that were processed using RNAscope, and visualized with confocal microscopy. FG labelled-positive neurons appear white, those expressing mRNA for somatostatin are green, while CRH or dynorphin mRNA positive neurons are in red (Figs. 6C and 6D, respectively). Not only did CeA FG labelled-positive neurons express mRNA associated with one of these phenotypes, a substantial subpopulation co-expressed mRNA for somatostatin and either CRH or dynorphin.

We quantified the number of retrogradely labeled CeA-LPB neurons displaying these phenotypes. Figure 6E depicts the proportion of CeA FG-positive neurons that expressed each phenotype, or that were double-labelled for two phenotypes, as median and 95% confidence intervals. Although separate sections were treated with primers either for somatostatin and CRH (n = 21 sections, 5 animals), or somatostatin and dynorphin (n = 23 sections, 5 animals), there was no statistically significant difference between the proportion of somatostatin labelled-positive neurons in these samples, therefore, we pooled data from both assays. We included in these analyses only those sections that contained a minimum of 20 FG-positive neurons, to control for variation in the intensity of FG deposition and uptake at the injection site. Dynorphin expressing neurons accounted for the largest fraction of these phenotypes. The median fraction of FG labelled-positive CeA-LPB neurons that express dynorphin was 0.42 (95% CI = 0.28 - 0.49), compared to a median of 0.26 for somatostatin (95% CI = 0.22 - 0.31), and a median of 0.09 for CRH (95% CI = 0.04 - 0.22). The median fraction of FG labeled neurons that expressed somatostatin and dynorphin was 0.15 (95% CI = 0.11 - 0.24), compared to a median of 0.03 (95% CI = 0 - 0.08) for somatostatin and CRH.

These findings indicate that of neurons comprising the inhibitory CeA-PB, almost half express dynorphin, and many of these neurons co-express somatostatin. A smaller population of neurons in this pathway express somatostatin alone and/or CRH. We did not probe for other neurochemical markers of CeA neurons, such as protein kinase C-δ or neurotensin (Luisa Torruella-Suárez et al., 2019; Wilson et al., 2019).

### Causality

To determine if the CeA-PB pathway plays a causal role in the regulation of pain sensation and the development of chronic pain, we tested the prediction that manipulating the CeA-PB pathway—pharmacologically or optogenetically—would result in changes in behavioral metrics of pain. Consistent with a causal role in chronic pain for this pathway, disinhibiting the parabrachial caused increases in pain metrics, whereas inhibiting the PB alleviated signs of chronic pain; Activating the CeA-LPB pathway reduced metrics of acute pain.

#### Pharmacological inhibition of PB

We first tested the prediction that inhibition of PB will suppress pain metrics in animals with CCI-Pain. To inhibit PB we infused muscimol, the GABA_A_ receptor agonist, or a saline-vehicle control into the LPB of CCI-ION injured rats (see Methods).

Figure 7B-C depicts the facial mechanical withdrawal thresholds for individual animals, as well as the median and 95% confidence intervals for each time point. Ipsilateral to the injury, left face thresholds changed across time points (Friedman’s F = 10.22; p = 0.0065), from a median of 34.8g at baseline (95% CI = 29.9 - 37.0g) to 20.47g after CCI (95% CI = 14.9 - 30.3g), 24.9g after saline (95% CI = 13.3 - 32.5g), and 34.8g after muscimol infusion (95% CI = 25.4 - 47.5g). Individual comparisons (after Dunn’s correction) revealed a difference (at α = 0.05) between baseline and CCI (p=0.05; Cohen’s *ds* = 2.87), but not between baseline and saline (p=0.2) or between baseline and muscimol (p>0.99). Right face (contralateral) mechanical withdrawal thresholds did not change across time points (Friedman’s F = 6.830; p = 0.07). Median mechanical withdrawal thresholds were 34.8g at baseline (95% CI = 32.5 - 34.8g), 24.2g after CCI (95% CI = 17.0 - 32.5g), 22.7g after saline (95% CI = 10.7 - 37.0g), and 32.5g after muscimol (95% CI = 27.2 - 37.0g).

**Figure 7.**
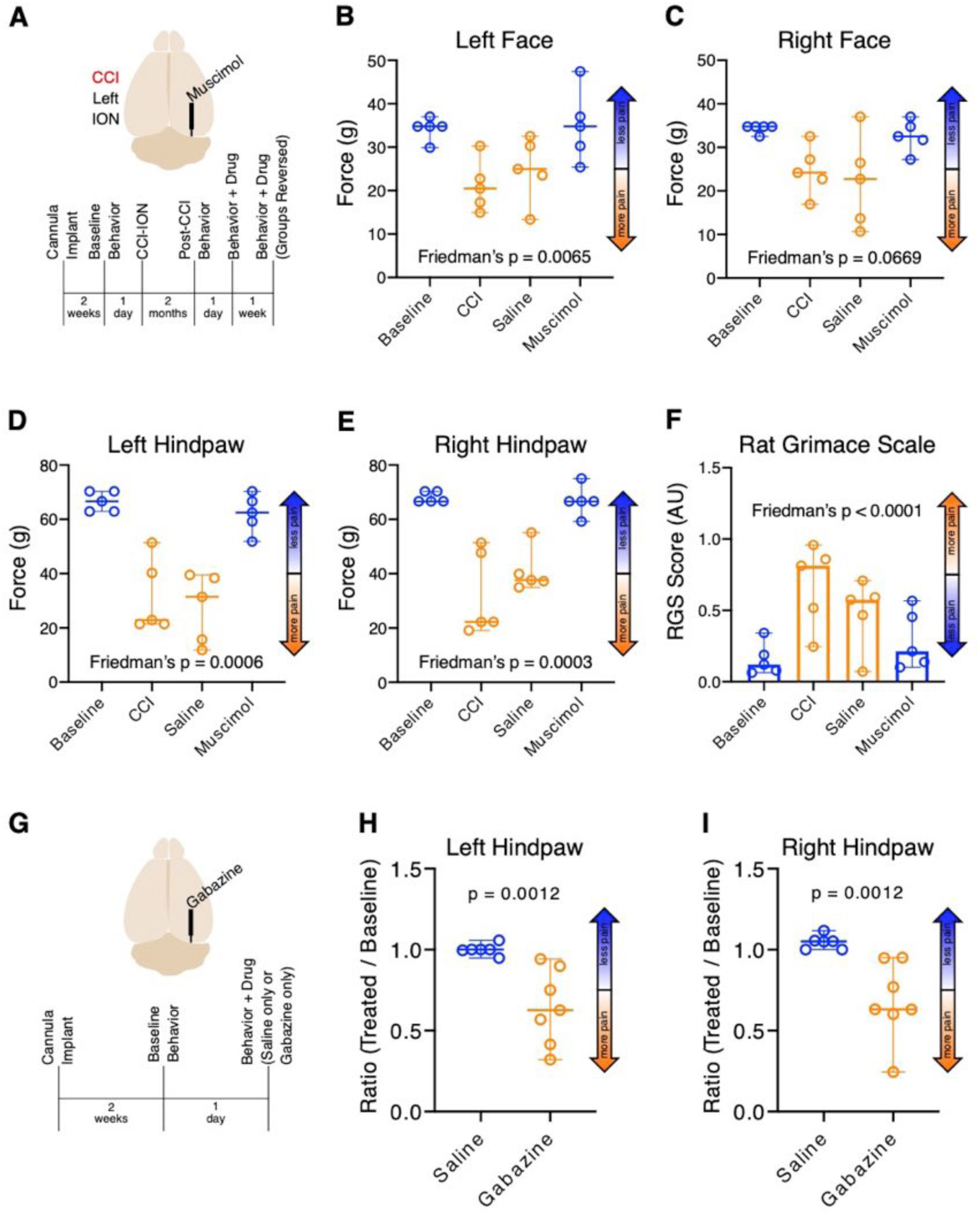
Pharmacologically inhibiting PB suppresses CCI-ION pain; In naïve animals, disinhibiting PB amplifies acute reflexive pain. Data are presented as medians and 95% confidence intervals. ***A***, Timeline of injury, cannula implantation, behavioral testing, and PB inhibition experimental design. ***B-F***, A single infusion of muscimol into the right LPB ameliorated the effect of CCI-ION on mechanical withdrawal thresholds in the left face (***B***) and both hindpaws (***D-E***), but not in the right face (***C***). Spontaneous pain RGS scores were also reduced by muscimol, but unaffected by saline (***F***). ***G***, Timeline of cannula implantation, behavioral testing, and PB disinhibition experimental design. ***H-I***, A single infusion of gabazine, but not saline, reduced hindpaw mechanical withdrawal thresholds both ipsilateral (***I***) and contralateral (***H***) to LPB cannulation.

Withdrawal thresholds to mechanical stimuli applied to both hindpaws were affected by muscimol but not saline (Fig. 7D-E). Left hindpaw mechanical withdrawal thresholds changed across time points (Friedman’s F = 12.60; p = 0.0006). Median mechanical withdrawal thresholds changed from 66.6g at baseline (95% CI = 62.9 - 70.3g) to 22.9g after CCI (95% CI = 21.4 - 51.4g), 31.4g after saline infusion (95% CI = 11.9 - 39.5g), and 62.5g after muscimol application (95% CI = 51.8 - 70.3g). Individual comparisons (using Dunn’s correction for multiple comparisons) revealed a difference between baseline and CCI (p=0.02; Cohen’s *ds* = 3.51), baseline and saline (p=0.0099; Cohen’s *ds* = 4.15), but not between baseline and muscimol (p>0.99; Cohen’s *ds* = 0.80).

We observed similar results on the contralateral hindpaw (Fig. 7E). Right hindpaw mechanical withdrawal thresholds changed across timepoints (Friedman’s F = 13.04; p = 0.0003). Median mechanical withdrawal thresholds changed from 66.6g at baseline (95% CI = 66.6 - 70.3g) to 22.2g after CCI (95% CI = 19.1 - 51.4g), 37.7g after saline application (95% CI = 34.9 - 55.1g), and 66.6g after muscimol infusion (95% CI = 59.2 - 75.0g). Individual comparisons (after Dunn’s correction) revealed a difference between baseline and CCI (p=0.007; Cohen’s *ds* = 3.19), but not between baseline and saline (p=0.06) or baseline and muscimol (p>0.99).

RGS scores, reflecting ongoing (“spontaneous”) pain, changed across time points (Friedman’s F = 14.04; p < 0.0001; Fig. 7F), from a median of 0.12 action units (AU) at baseline (95% CI = 0.06 - 0.34 AU) to 0.81 AU after CCI (95% CI = 0.25 - 0.96 AU). Post-CCI, infusion of saline caused a small decrease to 0.57 AU (95% CI = 0.07 - 0.71 AU), whereas infusion of muscimol caused a dramatic decrease to 0.21 AU (95% CI = 0.10 - 0.57 AU). Individual comparisons (using Dunn’s corrections) revealed a difference between baseline and CCI (p = 0.0007; Cohen’s *ds* = −2.35), but not between baseline and saline (p = 0.08), or baseline and muscimol (p = 0.4).

These findings indicate that pharmacological inhibition of PB attenuates both reflexive and ongoing signs of chronic pain.

#### Pharmacological disinhibition of PB

We tested also the corollary prediction, that disinhibition of PB will amplify behavioral metrics of pain. We did this by infusing gabazine, a GABA_A_ receptor antagonist, into LPB of naïve rats. Consistent with previous reports (Han et al., 2015; Barik et al., 2018; Chiang et al., 2019) this procedure appeared to be aversive to most animals, often resulting in either freezing or escape behaviors. This made it difficult for us to quantify pain-related behaviors in most of these animals.

In a subset of these animals we were able to evaluate hindpaw mechanical withdrawal thresholds (Fig. 7H-I). In the left hindpaw, the thresholds of gabazine treated animals (n=7) were reduced to 0.65 of baseline values (95% CI = 0.32 - 0.94), whereas ratios for saline controls (n=6) remained unchanged at 1.00 (95% CI = 0.95 - 1.06; Mann-Whitney *U* = 0; p = 0.0012; Cohen’s *ds* = 2.03). In the right hindpaw—ipsilateral to the gabazine injections—mechanical thresholds of gabazine treated animals were reduced to a median of 0.63 (95% CI = 0.24 - 0.95), whereas saline controls had a median ratio of 1.05 (95% CI = 1.00 - 1.12; Mann-Whitney *U* = 0; p = 0.0012; Cohen’s *ds* = 2.00). These findings suggest that unilateral disinhibition of PB results in reduced mechanical withdrawal thresholds to stimuli applied bilaterally to the hindpaws.

#### Optogenetics

Our findings that inhibiting PB attenuates chronic pain, and that CeA provides the major inhibitory input to PB, suggest that the inhibitory CeA-PB pathway is causally involved in regulating the experience of perception of pain and aversion. To test this hypothesis, we optogenetically excited CeA axon terminals in LPB while monitoring behavioral metrics of pain. We injected hChR2 viral constructs (see Methods) into the right CeA of wild-type or transgenic CRH-Cre rats, and implanted fiber optic cannulae over the right LPB. Although, the fluorescent reporter protein expressed by the viral construct can be observed in LPB at 6 weeks, expression appears to peak and stabilize around 8-9 weeks post-injection (similar to previous reports, Pomrenze et al., 2015). We used animals of both sexes, but did not have sufficient sample sizes to test for sex differences. To avoid potential confounds related to long-duration optogenetic stimulation (Tyssowski and Gray, 2019), we delivered short pulses of 470nm light (a single train of 5 light pulses, 4ms duration/pulse, 10Hz) only during application of mechanical or dynamic stimuli.

##### Mechanical withdrawal

We first tested the effects of activating CeA-LPB terminals in PB on mechanical allodynia in wild-type animals. Figure 8B-C depicts the hindpaw mechanical withdrawal thresholds for individual animals (n = 9) as well as the median and 95% confidence intervals for each time point. Left hindpaw thresholds increased from a median of 52.3g at baseline (95% CI = 52.3 - 53.2g) to 84.8g during optical stimulation (95% CI = 65.4 - 84.8g; Wilcoxon p = 0.004; Cohen’s *ds* = 2.49). Similarly, right hindpaw thresholds increased from a median of 52.3g at baseline (95% CI = 44.9 - 53.2g) to 72.8g during optical stimulation (95% CI = 53.2 - 84.8g; Wilcoxon p = 0.004; Cohen’s *ds* = 1.54).

**Figure 8.**
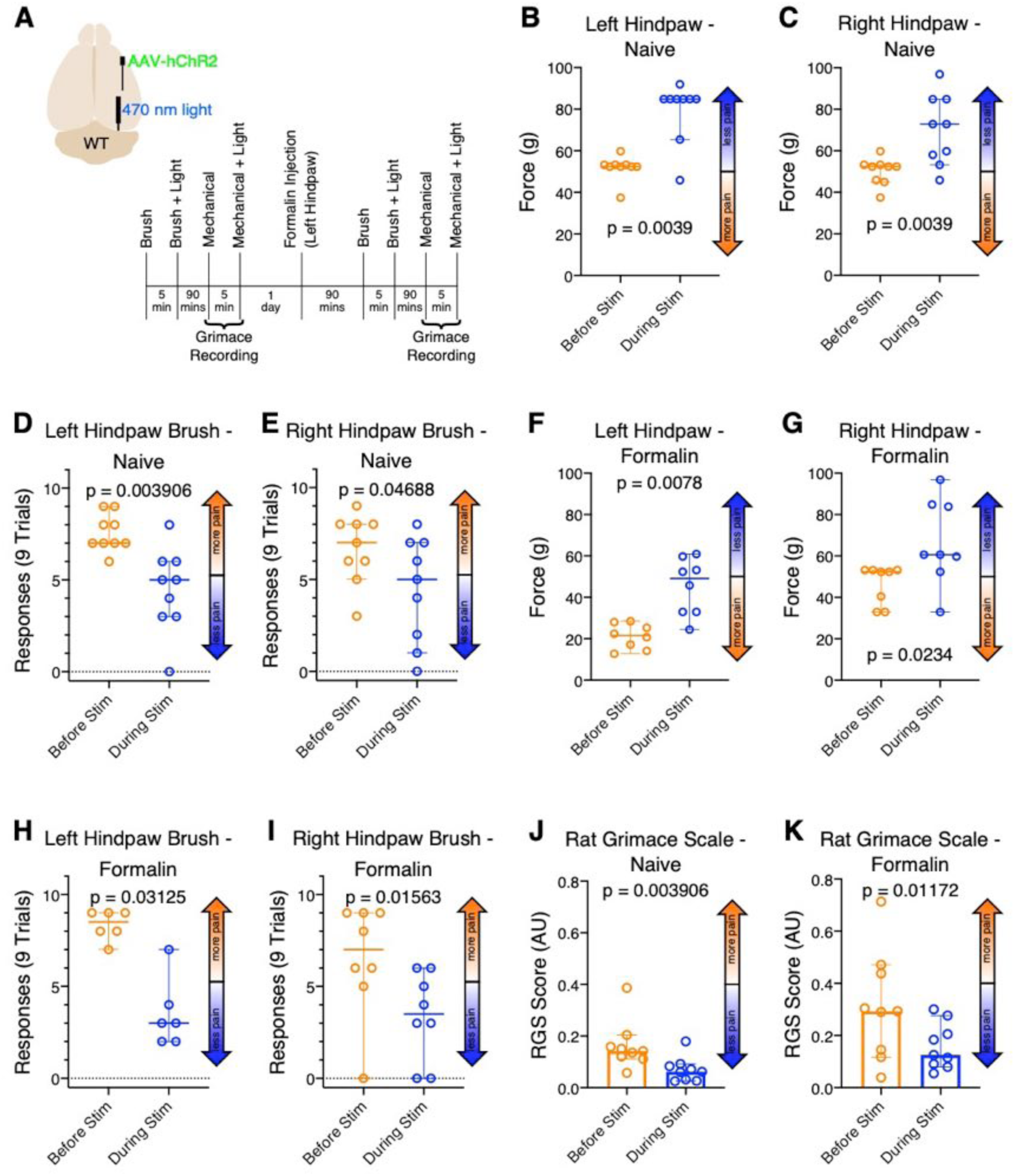
Optogenetically activating CeA-LPB terminals suppresses pain in the naïve animal and during acute formalin pain. ***A***, Locations of viral construct injection, fiber optic implantation, and experimental design. ***B-K***, Data are presented as medians and 95% confidence intervals. Activating right side CeA-LPB terminals using short pulses of light increased mechanical withdrawal thresholds in both hindpaws (***B-C***), and decreased both mechanical allodynia responses (***D-E***) as well as spontaneous pain RGS scores (***J***) in the naïve animal. During acute formalin pain, optogenetically activating CeA-LPB terminals also increased hindpaw mechanical withdrawal thresholds (***F-G***), and decreased both mechanical allodynia responses (***H-I***) as well as RGS scores (***K***).

We repeated this experiment in CRH-Cre animals, in which hChR2 is expressed only in the subset of GABAergic CeA neurons expressing CRH. Like in wild-type animals, left hindpaw mechanical withdrawal thresholds increased from a median of 53.2g at baseline (95% CI = 37.5 - 72.8g) to 84.8g during optical stimulation (95% CI = 52.3 - 96.8g; Wilcoxon p = 0.016; Cohen’s *d_s_* = 1.81; Fig. 9B). Similarly, right hindpaw thresholds increased from a median of 52.3g at baseline (95% CI = 37.5 - 60.6g) to 84.8g during optical stimulation (95% CI = 44.9 - 96.8g; Wilcoxon p = 0.016; Cohen’s *d_s_* = 1.68; Fig. 9C).

**Figure 9.**
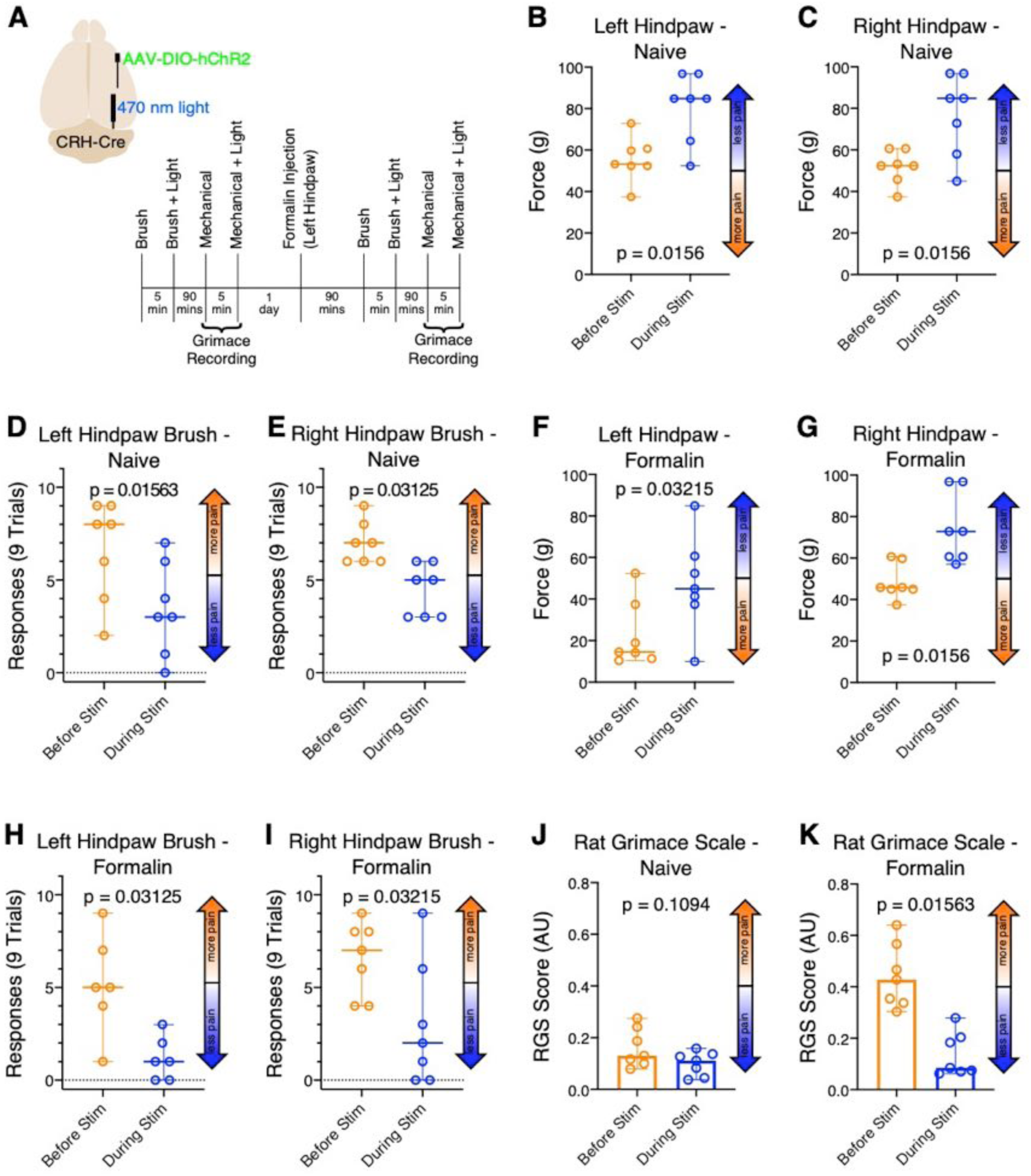
Optogenetically activating a subset of GABAergic CeA-LPB axon terminals in CRH-Cre rats suppresses pain in naïve animals, as well as during acute formalin pain. ***A***, Locations of viral construct injection, fiber optic implantation, and experimental design. ***B-K***, Data are presented as medians and 95% confidence intervals. Selectively activating CRH CeA terminals in LPB increased mechanical withdrawal thresholds in both hindpaws (***B-C***), and decreased mechanical allodynia responses (***D-E***), but did not affect spontaneous pain RGS scores (***J***) in the naïve animal. During acute formalin pain, optogenetically activating this subset of CeA-LPB axon terminals also increased hindpaw mechanical withdrawal thresholds(***F-G***), and decreased both mechanical allodynia responses (***H-I***) as well as RGS scores (***K***).

##### Dynamic mechanical allodynia

Activating CeA-LPB terminals in PB also reduced responses to the hindpaw brush test (see Methods). Figure 8D-E depicts the number of times each animal responded to the brush stimuli. In wild-type animals (n=9), left hindpaw brush responses decreased from a median of 7 of 9 stimuli at baseline (95% CI = 7 - 9 responses) to 5 responses during optical stimulation (95% CI = 3 - 6; Wilcoxon p = 0.004; Cohen’s *ds* = 1.75). Similarly, right hindpaw brush response decreased from a median of 7 responses at baseline (95% CI = 5 - 8 responses) to 5 responses during optical stimulation (95% CI = 1 - 7 responses; Wilcoxon p = 0.047; Cohen’s *ds* = 0.92).

In CRH-Cre animals (n=7), left hindpaw brush response decreased from a median of 8 responses out of 9 stimulus applications at baseline (95% CI = 2 - 9 responses) to 3 responses during optical stimulation (95% CI = 0 - 7 responses; Wilcoxon p = 0.016; Cohen’s *ds* = 1.21; Fig. 9D). Similarly, right hindpaw brush response decreased from a median of 7 responses at baseline (95% CI = 6 - 9 responses) to 5 responses during optical stimulation (95% CI = 3 - 6 responses; Wilcoxon p = 0.03; Cohen’s *ds* = 2.01; Fig. 9E).

##### Formalin-induced pain

To test the effects of activating the CeA-PB pathway on ongoing pain, we injected, one day after baseline testing, 5% formalin (50 µL) into the dorsal surface of the left hindpaw (contralateral to CeA virus injection and LPB cannula implantation). Left hindpaw mechanical withdrawal thresholds increased from a median of 21.6g after formalin exposure (95% CI = 12.7 - 28.5g) to 49.1g during optical stimulation (95% CI = 24.4 - 60.9g; Wilcoxon p = 0.008; Cohen’s *ds* = 2.29; Fig. 8F). Similarly, right hindpaw thresholds increased from a median of 52.3g after formalin exposure (95% CI = 33.0 - 53.2g) to 60.6g during optical stimulation (95% CI = 33.0 - 96.8g; Wilcoxon p = 0.02; Cohen’s *ds* = 1.25; Fig. 8G).

Similar effects of optogenetic activation occurred in CRH-Cre rats (n = 7; Fig. 9F-G). Left hindpaw withdrawal thresholds increased from a median of 14.5g after formalin exposure (95% CI = 10.4 - 52.3g) to 44.9g during optical stimulation (95% CI = 10.0 - 84.8g; Wilcoxon p = 0.03; Cohen’s *ds* = 1.25). Similarly, right hindpaw thresholds increased from a median of 45.8g after formalin exposure (95% CI = 37.5 - 60.6g) to 72.8g during optical stimulation (95% CI = 57.0 - 96.8g; Wilcoxon p = 0.016; Cohen’s *ds* = 1.91).

Optical stimulation of the CeA-PB pathway also attenuated dynamic mechanical allodynia evoked by the formalin injection. After formalin injection in the left hindpaw of wild-type animals (n = 6 animals), brush response to the same paw decreased from a median of 8.5 responses out of 9 stimuli (95% CI = 7 - 9 responses) to 3 responses during optical stimulation (95% CI = 2 - 7; Wilcoxon p = 0.0312; Cohen’s *ds* = 3.35; Fig. 8H). Similarly, right hindpaw brush response (n = 8 animals) decreased from a median of 7 responses after formalin exposure (95% CI = 0 - 9 responses) to 3.5 responses during optical stimulation (95% CI = 0 - 6; Wilcoxon p = 0.0156; Cohen’s *ds* = 1.14; Fig. 8I).

We obtained similar results in CRH-Cre animals (n=6; Fig. 9H-I). Left hindpaw brush responses decreased from a median of 5 out of 9 stimuli after formalin exposure (95% CI = 1 - 9 responses) to 1 response during optical stimulation (95% CI = 0 - 3 responses; Wilcoxon p = 0.03; Cohen’s *ds* = 1.91). Similarly, right hindpaw brush response (n = 7 animals) decreased from a median of 7 of 9 responses after formalin exposure (95% CI = 4 - 9 responses) to 2 responses during optical stimulation (95% CI = 0 - 9 responses; Wilcoxon p = 0.03; Cohen’s *ds* = 1.29).

##### Grimace

Stimulation of the amygdalo-parabrachial also attenuated grimace scores in rats with formalin-induced pain (Fig. 8J-K). In naïve animals (n=9), activating CeA-LPB reduced median RGS scores from 0.14 action units (AU) (95% CI = 0.11 - 0.20 AU) at baseline to 0.06 AU (95% CI = 0.03 - 0.09 AU) during light stimulation (Wilcoxon p = 0.004; Cohen’s *ds* = 1.26). After formalin injection, activation of CeA-PB reduced median RGS scores from 0.29 AU (95% CI = 0.12 - 0.47 AU) during acute formalin pain to 0.12 AU (95% CI = 0.08 - 0.28 AU) during optical stimulation (Wilcoxon p = 0.01; Cohen’s *ds* = 0.96).

We replicated these results in CRH-Cre animals (n = 7; Fig. 9J-K). In naïve animals, activating CeA-LPB reduced median RGS scores only nominally, from 0.13 action units(AU) (95% CI = 0.08 - 0.27 AU) at baseline to 0.11 AU (95% CI = 0.04 - 0.16 AU) during light stimulation (Wilcoxon p = 0.11). However, after formalin injection, activation of CeA-LPB reduced median RGS scores from 0.43 AU (95% CI = 0.30 - 0.64 AU) during acute formalin pain to 0.08 AU (95% CI = 0.06 - 0.28 AU) during optical stimulation (Wilcoxon p = 0.016; Cohen’s *ds* = 2.86).

Taken together, these findings suggest that optogenetic activation of the CeA-PB pathway inhibits LPB neurons, resulting in attenuation of both acute and ongoing pain-related behaviors.

## Discussion

### Amplified activity of PB neurons

As detailed in our Introduction, the parabrachial complex (PB)—and, in particular, its lateral division (LPB)—is a critical nexus for processing nociception and for the perception of both sensory and affective components of pain. Growing evidence supports the notion that these pain related functions are part of a larger role of PB in mediating aversion (see Introduction).

Here, we report that a mouse model of chronic, neuropathic pain—constriction injury of the infraorbital nerve (CCI-ION)—results in lasting amplification of activity in LPB neurons. This amplification was expressed as increased spontaneous activity, and larger magnitude responses to noxious stimuli. Particularly striking was the increase in after-discharges, responses that far outlast stimuli. After-discharges have been described in other brain regions after chronic pain (Woolf and King, 1987; Herrero et al., 2000). Their durations, and the proportion of neurons that express them, is dramatically increased in chronic pain (Palecek et al., 1992; Laird and Bennett, 1993). After-discharges are also key in “windup” – the progressively increased excitability of spinal neurons that occurs in central sensitization to pain (Morisset and Nagy, 2000). After-discharges may increase pain perception by enhancing the transfer of nociceptive responses to downstream structures (Morisset and Nagy, 1998). There is compelling evidence that after-discharges are causally related to the presence of chronic pain (Laird and Bennett, 1993; Asada et al., 1996). For example, we have demonstrated that the incidence and duration of after-discharges in spinal neurons increases significantly in animals with chronic, neuropathic pain. Importantly, suppressing after-discharges significantly lessens hyperalgesia in experimental animals (Okubo et al., 2013).

The current finding in mice is similar to that in our previous study in rats, in which we reported a dramatic increase in after-discharges of PB neurons after CCI-ION (Uddin et al., 2018). However, contrary to our current findings, in the rat CCI-ION was not associated with changes in spontaneous firing rates of PB neurons, or in their responses during stimulus application. We do not yet know if these species differences reflect methodological differences, or more fundamental biological distinctions.

### Disinhibition of PB neurons

The amplification of PB neuronal activity was not related to changes in passive membrane properties, as neither the input resistance nor the resting membrane potential of PB neurons was affected by CCI-ION. What did change dramatically was the inhibitory input to these neurons. The frequency of miniature inhibitory postsynaptic currents (mIPSCs) was reduced by half, whereas the amplitude of mIPSCs remained unchanged, suggesting a reduction in presynaptic inhibitory inputs to PB neurons. That reduced inhibition in PB is related to chronic pain is consistent with findings in other CNS regions, where similar dis-inhibitory mechanisms are causally related to chronic pain (reviewed in Prescott, 2015; Todd, 2015). Reduced inhibition may directly result in the increased spontaneous and evoked responses in mouse PB neurons in chronic pain, as reported here.

It remains to be determined whether disinhibition gives rise to the pronounced after-discharges in PB neurons of rodents with chronic pain. In other neuronal populations, after-discharges appear to be mediated by interactions between synaptic inhibition, NMDA receptors and potassium currents (Traub et al., 1995; Sotgiu and Biella, 2002; Drew et al., 2004). Whether similar biophysics governs that pathophysiology of after-discharges in PB during chronic pain remains to be determined.

### Source of PB inhibition

We find that a relatively modest proportion of neurons (12% in rats) in PB are GABAergic. This is consistent with previous reports of a population of inhibitory neurons in the rodent PB (Chiang and Ross, 2017; Chen et al., 2017; Geerling et al., 2017; Chiang et al., 2019). Here, we focused on the role of extrinsic GABAergic inputs to PB.

### The CeA-PB pathway

We identified several extrinsic sources of GABAergic inputs to PB, including the zona incerta, hypothalamus, and, most prominent, the central nucleus of the amygdala (CeA). Our results show that this inhibitory CeA-PB amygdalo-parabrachial pathway plays a key role in regulating pain-related activity involving PB.

It is well established that CeA is a central node for pain processing, integrating nociception, contexts, affect, and memory to guide behaviors related to acute and chronic pain (Davis, 1994; Zald, 2003; Veinante et al., 2013; Neugebauer, 2015; Woodhams et al., 2017; Wilson et al., 2019).

We describe a dense, GABAergic pathway, originating from CeA and innervating PB. This pathway exists in both rats and mice, and provides potent inhibition to PB neurons. We show that, in both rodent species, this CeA-PB amygdalo-parabrachial inhibitory pathway regulates responses of PB neurons to noxious stimuli and behaviors evoked by these stimuli. We also demonstrate that activating this inhibitory pathway ameliorates signs and of chronic pain, and that suppressing this pathway in naïve animals promotes pain behaviors. These findings support a causal role for the CeA-PB amygdalo-parabrachial pathway in both acute and chronic pain.

Our findings are consistent with prior anatomical reports that made reference to a direct projection from CeA to PB (Jia et al., 1994; Sun et al., 1994; Chieng et al., 2006; Pomrenze et al., 2015). Jia et al (2005) provided a more extensive description of afferents originating from CeA and terminating in the lateral PB, where they form symmetrical—presumably inhibitory—synapses. More recently, Luisa Torruella-Suárez et al (2019) reported that inhibitory, neurotensin-expressing CeA neurons densely innervate and inhibit PB, and that stimulation of these projections promotes consumption of ethanol and palatable fluids.

The GABAergic projections from CeA to PB complement the more extensively described reciprocal projections from PB to CeA (Saper and Loewy, 1980; Bernard et al., 1993; Sarhan et al., 2005), that are involved in mediating pain-related behaviors (Han et al., 2015). This PB->CeA pathway is a glutamatergic, excitatory one (Neugebauer, 2015), balanced by the inhibitory reciprocal pathway from CeA to PB described here. At least some PB neurons that project to CeA appear to receive inhibitory inputs from CeA, consistent with direct connections between individual CeA->PB and PB->CeA neurons (Jia et al., 2005). PB projections target both somatostatin and CRH neurons in specific subdivisions of CeA; in a model of chronic pain, the strengths of these projections may increase or decrease, depending on the neuronal population targeted in CeA (Li and Sheets, 2020). How these reciprocal pathways influence each other remains to be determined. We discuss several possibilities below.

Consistent with previous reports that essentially all CeA neurons are inhibitory (Ehrlich et al., 2009; Janak and Tye, 2015), we find that CeA neurons that project to PB are GABAergic. CeA neurons are functionally and neurochemically heterogenous (Janak and Tye, 2015; Wilson et al., 2019). We find that the GABAergic CeA neurons involved in the CeA-PB pathway are also heterogeneous, and include neurons that express dynorphin (42% of these projections neurons), somatostatin (26%) and CRH (9%), with some expressing more than one of these substances. We did not probe for other neurochemical phenotypes expressed in CeA, such as PKCδ (Wilson et al., 2019) or neurotensin (Luisa Torruella-Suárez et al., 2019). How these diverse populations of CeA-PB neurons regulate nociception and pain remains to be fully elucidated.

At least one of these populations—the somatostatin expressing neurons—has been directly implicated in these functions, as they are inhibited by nerve injury, and activating them attenuates mechanical allodynia (Wilson et al., 2019). These somatostatin CeA neurons are involved also in responses to threat-related sensory cues (Yu et al., 2016; Fadok et al., 2017). Carrasquillo and colleagues (Wilson et al., 2019) demonstrated that two populations of CeA neurons are involved in opposing functions in pain regulation. PKCδ expressing neurons are involved in functions that promote pain (pro-nociceptive). Whereas somatostatin-CeA neurons are involved in opposing, anti-nociceptive functions. Our present results suggest that the anti-nociceptive function of these somatostatin-CeA neurons is accomplished by their direct, inhibitory action upon PB neurons.

The particularly large population of CeA-PB neurons that express dynorphin is also of significance because the dynorphin receptor, the kappa-opioid receptor, is highly expressed in PB (Unterwald et al., 1991; Yasuda et al., 1993; Mansour et al., 1994). How dynorphin regulates PB neuronal activity is still unknown. By analogy to other CNS structures, where these functions are better understood in the context of pain (Bie and Pan, 2003; Wang et al., 2018; Navratilova et al., 2019), we speculate that dynorphin, acting on kappa opioid receptors, may directly inhibit PB neurons thorough postsynaptic receptors, as well as regulate GABA release presynaptically. We are currently studying these mechanisms.

Taken together, our findings demonstrate a hitherto unknown pathway of pain regulation, involving two key structures associated primarily with the affective aspects of chronic pain, the parabrachial complex and the amygdala. Because the affective component of chronic pain is the component most directly associated with patients’ suffering, and is the component most resistant to treatment (Han et al., 2015; Neugebauer, 2015; Price et al., 2018; Corder et al., 2019), the identification of a novel pathway mediating this function may be directly relevant to planning novel therapies for this devastating condition.

## Acknowledgements

We are grateful to Dr. Adam Puche for sharing his histological expertise, and to Andrew Furman for sharing his Matlab routines. Research reported in this publication was supported by the National Institute of Neurological Disorders and Stroke of the National Institutes of Health grants R01NS099245 and R01NS069568. The content is solely the responsibility of the authors and does not necessarily represent the official views of the National Institutes of Health. The funding sources had no role in study design; the collection, analysis and interpretation of data; the writing of the report; or in the decision to submit the article for publication.

